# A droplet microfluidic-based platform for enhanced DNA delivery in non-model organisms

**DOI:** 10.64898/2026.04.30.721591

**Authors:** Aaron Stibelman, Alyse Tran, James Chappell, Yousif Shamoo

## Abstract

Expanding genetic engineering beyond model microorganisms is critical to unlocking novel applications in biotechnology, yet the low efficiency of DNA delivery methods like conjugation, remains a major bottleneck in non-model and environmental microbes. Here, we present an automated, high-throughput droplet microfluidic platform that enhances conjugation by encapsulating donor and recipient microbes in picoliter-scale water-in-oil microdroplets, stabilizing cell-cell contact and DNA transfer. Optimization of incubation time, donor to recipient ratio, and plasmid type yielded over a 100-fold increase in conjugation efficiency compared to conventional methods and enabled delivery of complex DNA libraries in low reaction volumes, demonstrating scalability for pooled plasmid library delivery. We further utilized a synthetic biology circuit for donor removal within microdroplets without antibiotic selection, eliminating the need for host-specific selection markers or engineered auxotrophs. When applied to a soil microbial community, this platform improved community-level conjugation, preserving microbial diversity and enabling the identification of genetically accessible chassis. Collectively, this platform establishes a scalable, generalizable solution for high throughput DNA delivery in previously inaccessible microbial hosts.

**Graphical abstract:** 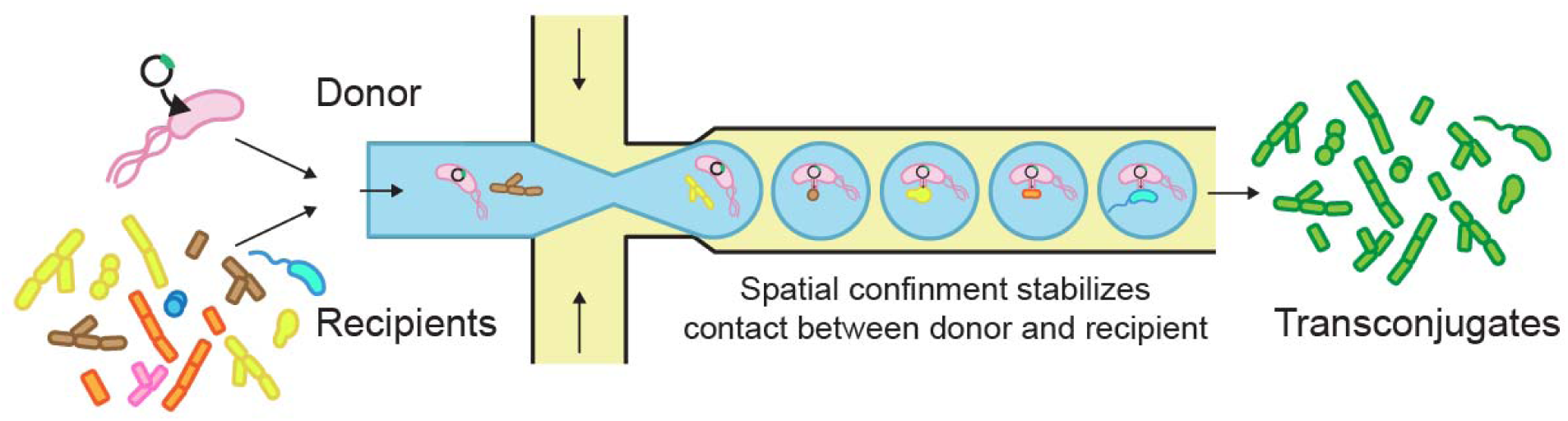

## Introduction

The expansion of synthetic biology beyond traditional model organisms is critical for developing improved microbial chassis with enhanced performance and functionality. Historically, the field has relied on a limited set of genetically tractable hosts, most notably *Escherichia coli*, which has enabled the rapid development of genetic tools supporting programmable genetic circuits, metabolic pathway engineering for sustainable production, and cell-based biosensing technologies (1–3). Despite these advances, reliance on a narrow set of model systems often lacks the environmental robustness and metabolic diversity required for many industrial and environmental applications. In contrast, non-model and environmental microbes offer a broad range of functional traits that make them attractive for industrial and environmental applications. Many exhibit inherent resistance to environmental stresses, including fluctuations in temperature, pH, and nutrient availability, enabling robust performance under variable field deployable conditions (4). Non-model organisms also often possess unique metabolic capabilities that could support more sustainable bioproduction strategies, such as the utilization of inexpensive or unconventional carbon sources (5). Furthermore, many environmental microbes are prolific producers of secondary metabolites relevant for human health, including antibiotics, anticancer compounds, and other bioactive molecules (6).

Despite these potential benefits, non-model microbes are often recalcitrant to genetic manipulation, especially the introduction of DNA, preventing downstream engineering. While chemical transformation and electroporation have been extensively optimized in model organisms (7), their effectiveness across diverse taxa is limited by variability in cell envelope architecture (8). As a result, these methods generally require extensive species-specific optimization (9). Additionally, exogenous DNA must evade host defense mechanisms such as restriction-modification systems, which frequently require organism-specific countermeasures (10). Delivery of DNA by conjugation provides a potential alternative, which relies upon dedicated cellular machinery to mediate DNA transfer between a donor and recipient cell. This process involves direct, contact-dependent transfer via Type IV secretion systems, delivering DNA as a single-stranded intermediate that is subsequently replicated within the recipient (11, 12). This enables incorporation of host-specific modifications, reducing recognition by host defense systems (13). Additionally, studies have shown that conjugative plasmids are often able to transfer to a surprisingly broad host range (14–16), potentially reducing the need for strain-specific approaches. Nevertheless, conventional conjugation workflows are limited by low transfer efficiencies, often several orders of magnitude lower in non-model organisms (17, 18), and low throughput due to their reliance on stochastic, contact dependent donor-recipient encounters, constraining their utility for large scale genetic engineering. These limitations in conventional conjugation workflows are particularly restrictive for emerging applications that require conjugation at scale, including pooled CRISPR-based perturbation screens and genetic manipulation of complex microbial communities.

Droplet microfluidic-based systems offer a promising strategy to massively multiplex reactions in high-throughput and increase the efficiencies of reactions by providing controlled, spatially confined microcellular environments that promote sustained cell-cell interactions (19). In these systems, aqueous solutions are partitioned into discrete picoliter-volume microdroplets with diameters ranging from tens to hundreds of micrometers within an immiscible oil phase, generating highly uniform and isolated compartments. Cells can be encapsulated individually or in small consortia within microdroplets, spatially confining them. In contrast to bulk or plate-based systems, where interactions arise from stochastic cell-cell encounters, microdroplet confinement maintains cells in close and sustained proximity. In community contexts, this compartmentalization also reduces interspecies competition observed in batch workflows by isolating cells into independent microenvironments, supporting more equitable species persistence. This spatial confinement has been broadly leveraged to enhance a range of proximity-dependent biological processes, including quorum sensing (20), metabolic cross-feeding (21), and microbial competition (22). By stabilizing cell-cell contact, microdroplet confinement also enables efficient contact-dependent processes such as conjugation. For example, recent work demonstrated intragenic conjugation between strains of *Bacillus subtilis* (23). Additionally, the automated, high-throughput nature of droplet microfluidics reduces the labor and time required to generate large libraries of transconjugates. Despite these advances, the application of droplet microfluidic platforms to perform cross-species conjugation into non-model organisms remains largely unexplored.

Here, we present a high-throughput droplet microfluidic platform to enhance conjugation-based DNA delivery into non-model soil microbes and microbial communities. We initially focused on *Streptomyces venezuelae*, a filamentous soil bacterium that produces clinically relevant secondary metabolites (24). Using *S. venezuelae* as a recipient, we systematically evaluated key parameters influencing conjugation efficiency, including incubation time, donor to recipient ratio, and plasmid type, achieving more than a 100-fold improvement compared to conventional plate-based methods, and is compatible with the introduction of large DNA plasmid libraries. We next established a synthetic biology circuit to allow for removal of donor cells in microdroplets, providing a donor removal mechanism that does not require host-specific antibiotic resistance or engineered auxotrophs, enabling deployment across diverse microbial hosts and communities. Finally, we demonstrate the platform’s applicability to enhance DNA transfer to a soil microbial community, showing roughly 50% conjugation efficiency across a group of tested isolates, compared to undetectable levels of conjugation using conventional methods. This platform establishes a versatile framework for high-throughput DNA delivery, enabling functional genomic studies and the construction of genetic libraries in previously challenging organisms.

## Results

### Co-encapsulation in microdroplets increases conjugation efficiency between *S. venezuelae* and an *E. coli* donor

To overcome the low efficiency of interspecies DNA transfer into bacteria, we developed a microdroplet-based microfluidic platform designed to physically constrain donor and recipient cells together, stabilizing the cell-cell interactions required for conjugation. We hypothesized that increasing and stabilizing cell-cell contact and more precise donor to recipient ratios, using microdroplet-confinement, would increase conjugation efficiencies (Figure 1a). Initially, microdroplet-based conjugation was carried out using *Escherichia coli* WM6029 as the donor strain (D) and *Streptomyces venezuelae* ATCC 19207 as a model recipient strain (R). The *E. coli* donor is auxotrophic for diaminopimelic acid (DAP), enabling donor counter selection, and is methylase negative to better accommodate the methyl-specific restriction system in *Streptomyces* (*25*). The donor was transformed with a conjugative, high copy number, *Streptomyces* compatible vector, pL97-R10, expressing mCherry and conferring resistance to apramycin.

**Figure 1.**
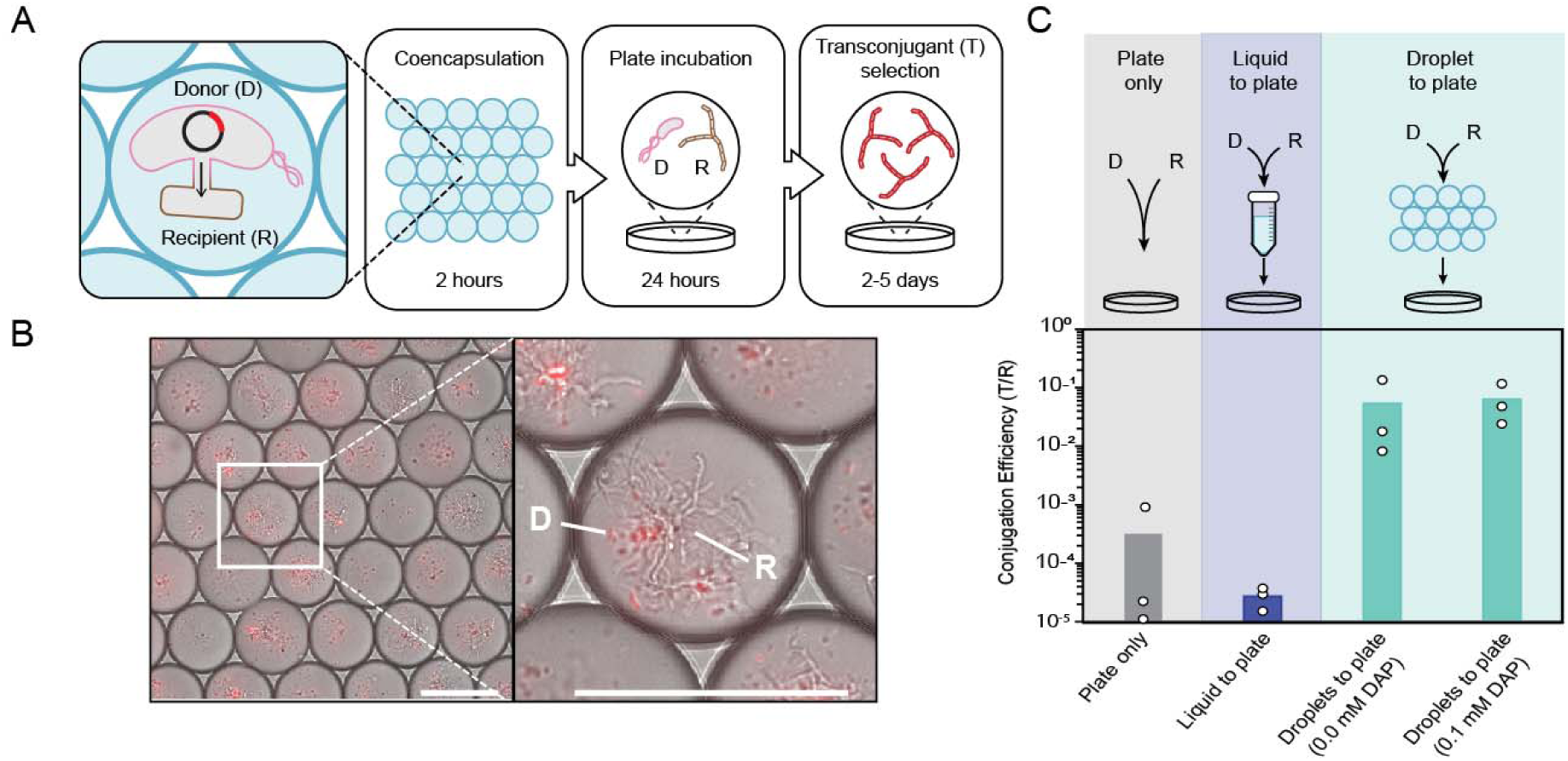
Microdroplet-based conjugation enhances conjugation efficiency compared to conventional methods. (A) Schematic of the microdroplet-based conjugation workflow. Donor (D) and recipient (R) cells are initially co-encapsulated within picoliter microdroplets for enhanced cell contact, then decapsulated, plated, and subsequently selected to recover transconjugants (T). (B) Microscopy image showing co-encapsulated donor *E. coli cells* containing an mCherry-expressing conjugative plasmid and recipient *S. venezuelae* mycelium within microdroplets (1:500 ratio). Scale bar: 100 µm. (C) Comparison of conjugation efficiencies across three methods: plate only incubation, liquid to plate incubation, and microdroplet to plate incubation. Conjugation efficiencies are calculated by normalizing transconjugant cells over recipient cells. For the microdroplet to plate approach, in droplet incubations were performed in the presence and absence of 0.1 mM DAP. Bars are the mean of three biological replicates.

To test whether the microdroplet-based approach enhanced conjugation efficiencies, we co-encapsulated *S. venezuelae* spores and stationary-phase donor cells at an average recipient-to-donor ratio of approximately 1:500 within each microdroplet. Since donor cell growth is contingent on the presence of DAP, we performed conjugations both with and without DAP to assess whether donor growth influenced conjugation efficiencies. Following 24-hour incubation within microdroplets, both donor and recipient cells were observed to coexist within the droplets, indicating successful co-culture under confined conditions (Figure 1b). Microdroplets were then plated onto nonselective solid media for an additional 24 hours prior to antibiotic selection to isolate transconjugants. Control conjugations were also performed in parallel using conventional methods, by mixing the same donor and recipient cell ratio on solid media plates (plate only) or incubating cells in liquid media for 24-hours prior to plating (liquid to plate) (Figure 1c). Following growth on plates, conjugation efficiency was determined by normalizing the number of recovered transconjugates to the total number of plated recipients. These results showed that microdroplet-based conjugation efficiency was 100-fold and 1000-fold greater than plate and liquid-based conjugations respectively. Notably, conjugation efficiency within microdroplets was indistinguishable between DAP supplemented and unsupplemented conditions (Figure 1c) suggesting that donor growth did not impact conjugation efficiency. We also compared the absolute number of recovered recipients to the absolute number of recovered transconjugants. While hundreds of transconjugants per event were observed when cells were incubated only on plates, microdroplet incubation resulted in the recovery of 14,000 to 80,000 transconjugants (Figure S1). To confirm that transconjugants contained the mCherry-expressing plasmid, five randomly selected transconjugants were analyzed by fluorescence microscopy and all confirmed to express mCherry (Figure S2). Taken together, these results demonstrate that microdroplet confinement during conjugation can increase both the conjugation efficiency and the number of transconjugates compared to conventional methods.

### Microdroplet-based conjugation efficiency is donor and time dependent

To further evaluate and optimize microdroplet-based conjugation, we systematically investigated the effects of donor to recipient cell ratio and incubation time on conjugation efficiency using three distinct plasmid types: the previously characterized pL97, the medium-copy plasmid pSG5, and a VWB phage-based integrative plasmid, pVWB (26). Conjugations were performed across a range of donor to recipient ratios and incubation times (2, 24, or 48 hours). To maintain a constant donor to recipient ratio throughout incubation, microdroplet-based conjugations were performed in the absence of DAP, preventing donor replication and ensuring that observed differences reflected incubation time and donor loading rather than donor overgrowth.

For pL97, we initially tested lower donor to recipient ratios (5:1, 50:1, and 500:1) and observed that conjugation efficiency increased with higher donor amounts, reaching a maximum at 1:500. Increasing the donor ratio beyond this point (5,000:1 and 25,000:1) did not further improve efficiency, indicating a plateau in transfer efficiency once a threshold donor density is reached (Figure 2A, S3). Consistent with this, conjugation efficiency depended on donor to recipient ratio at lower donor densities, but it became insensitive to donor ratio and incubation time above this, suggesting that under microdroplet confinement, pL97 transfer is robust once a minimal donor density is achieved. Based on this result, subsequent experiments were performed using higher donor to recipient ratios (500:1, 5,000:1, and 25,000:1) to ensure that donor availability was not limited.

**Figure 2.**
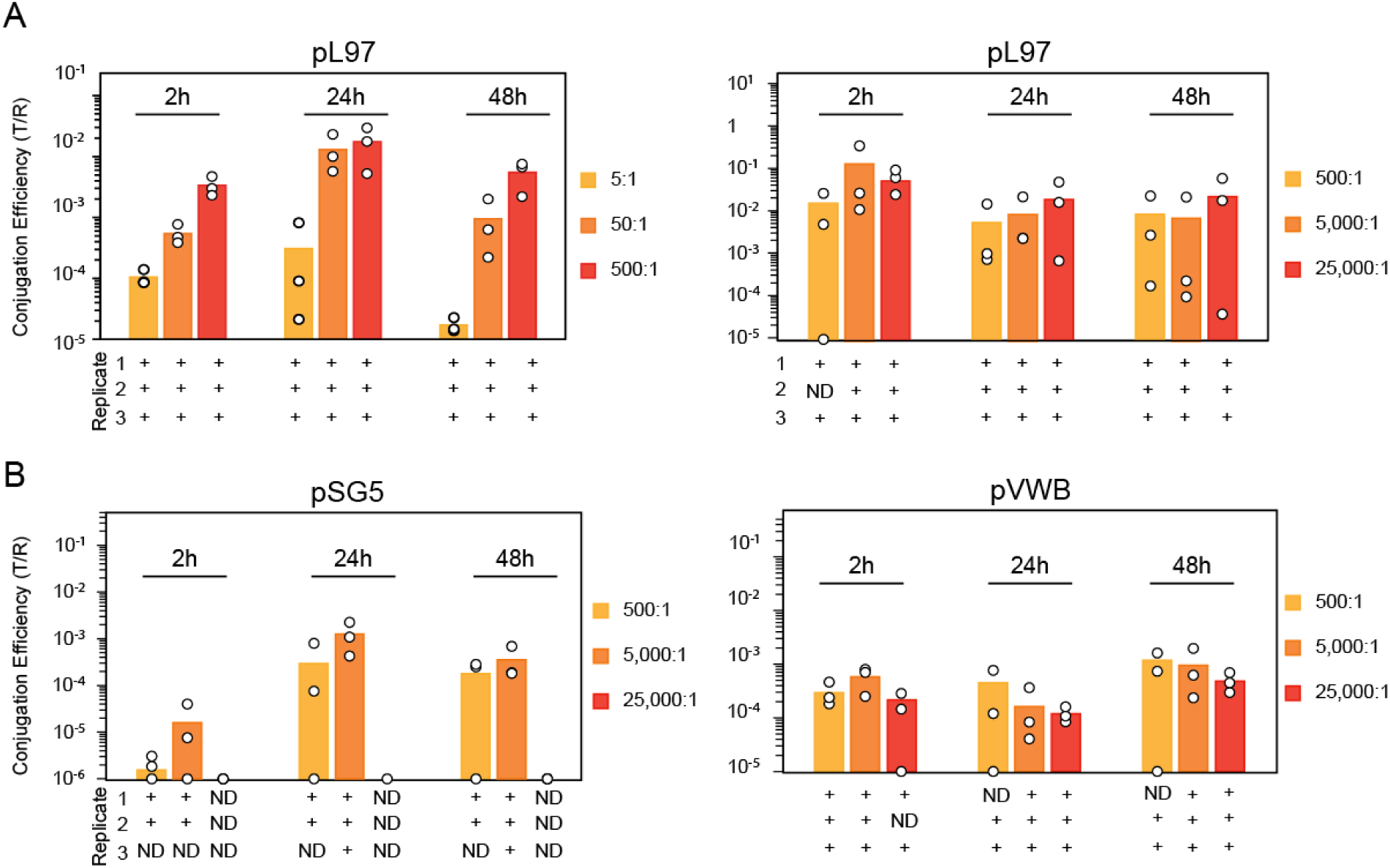
Conjugation efficiency using different plasmids can be optimized readily by varying incubation time. (a) Conjugation efficiency of the high-copy plasmid pL97 measured across incubation times (2, 24, 48 hours) and increasing donor:recipient ratios (5:1, 50:1, 500:1). Higher donor ratios led to increased conjugation efficiency, prompting further testing at elevated ratios (500:1, 5,000:1, 25,000:1), where efficiencies plateaued. (b) Conjugation efficiency of medium-copy (pSG5) and integrating (pVWB) plasmids measured across incubation times (2, 24, 48 hours) at donor:recipient ratios of 500:1, 5,000:1, and 25,000:1. Conjugations were carried out by co-encapsulating donor *E. coli* cells containing plasmids with recipient *S. venezuelae* cells. Conjugation efficiency was calculated by normalizing transconjugant cells (T) to recipient cells. Replicates below the detection limit (not detected, ND) were excluded from the average. Points represent individual biological replicates, and bars indicate the mean of 2–3 biological replicates.

In contrast, pSG5 conjugation exhibited a clear dependence on incubation time (Figure 2B). Increasing incubation duration led to higher conjugation efficiencies. Donor to recipient ratio did not significantly affect conjugation efficiency up to a ratio of 1:5,000; however, at the highest donor load (1:25,000) transconjugants were not recovered. This loss of detectable transfer coincided with a pronounced decrease in the total number of recovered recipients as donor density increased, suggesting that excessive donor loading may negatively impact recipient viability or growth rather than conjugation efficiency (Figure S4).

For the VWB integrative plasmid, higher incubation time increased conjugation efficiency. This is consistent with the additional requirement for site-specific recombination and chromosomal integration following DNA transfer. Donor to recipient ratio had comparatively little effect within the tested range, indicating that prolonged donor and recipient contact, rather than increased donor abundance, was the dominant factor for successful integration.

Together, these results demonstrate that optimal microdroplet conjugation conditions are strongly plasmid dependent. While pL97 transfer is relatively insensitive to both donor density and incubation time, pSG5 conjugation is limited by excessive donor loading, and VWB-mediated integration benefits from extended incubation. The lower efficiency observed for the integrative plasmid is consistent with the requirement for two sequential processes, conjugative transfer followed by chromosomal integration. These findings highlight the importance of optimizing microdroplet-based conjugation parameters to plasmid architecture and transfer mechanisms rather than applying a single set of conditions universally.

### Droplet microfluidics preserves plasmid library diversity during conjugation

To determine whether microdroplet-based conjugation could support the transfer of plasmid libraries, we constructed barcoded plasmid libraries in which each plasmid carries a distinct sequence that can be used to barcode a single conjugation event. This was achieved by cloning a DNA barcode composed of ten degenerate nucleotides into a VWB integrative plasmid. We hypothesized that by introducing different barcoded plasmids into the donor population, each successful conjugation and chromosomal integration event could be tracked through barcode sequencing, allowing us to track the number of independent conjugation events, and the composition and diversity of the library.

Three independent DNA plasmid libraries were generated, transformed directly into the donor strain, and subsequently conjugated using the microdroplet-based conjugation workflow (Figure 3a). Conjugations were performed using 100 µL of microdroplets, a volume comparable to that typically used in conventional plate-based conjugation approaches. Following incubation and recovery, genomic DNA was extracted from the resulting transconjugant populations, and barcode sequences were amplified directly from recipient genomes. Because the VWB vector integrates site-specifically into the chromosome (27), only plasmids that successfully transferred and integrated into recipients were detected. This approach ensured that each recovered barcode corresponded to an integration event while eliminating false positives arising from residual donor cells or nonintegrated DNA.

**Figure 3.**
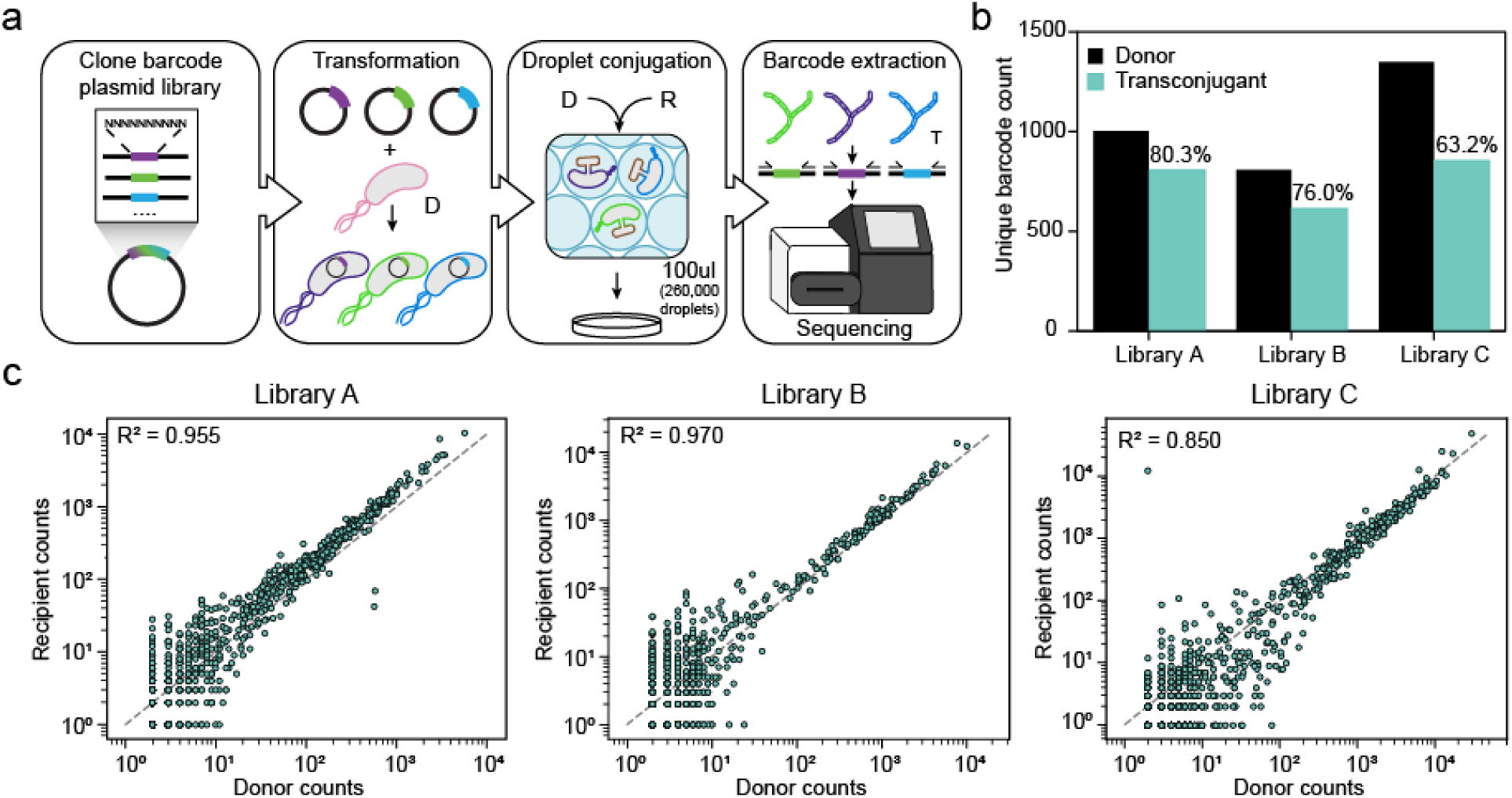
Droplet microfluidics maintains plasmid library diversity for conjugation between *E. coli* and *S. venezuelae* cells. (a) Schematic of the workflow. Three distinct barcoded pVWB-based plasmid libraries (A to C) were independently conjugated into recipient cells. Following conjugation, genomic DNA was extracted and barcodes were amplified directly from transconjugant genomes, ensuring that only integrated plasmids were detected and eliminating false positives from donor cells or non-integrated plasmids. (b) Number of unique barcodes detected by sequencing in each starting donor library and in the corresponding transconjugant populations following conjugation. (c) Scatter plots comparing the abundance of each unique barcode in the donor library versus the corresponding transconjugant population for each of the three libraries. Each point represents a single barcode.

We first quantified the number of unique barcodes present in each donor input library and compared these values to the number recovered in the corresponding transconjugant populations (Figure 3b). Because each barcode uniquely labels a plasmid variant, the number of distinct barcodes should represent the diversity of the library before and after transfer. Across all three libraries, microdroplet-based conjugation preserved a substantial fraction of the initial barcode identity, with 63.2%-80.3% of unique barcodes recovered following transfer. Although a reduction in total barcode count was observed following conjugation, the majority of uniquely represented variants were maintained within the transconjugant populations, indicating that microdroplet confinement does not impose a strong bottleneck on diversity under the conditions tested. Importantly, because microdroplet generation is readily scalable, total reaction volume can be increased beyond 100 µL to further improve library coverage and support the transfer of larger plasmid pools when required.

To further assess how well relative barcode abundances were maintained through conjugation and integration, we compared the abundance of each barcode in the donor libraries to its abundance in the corresponding transconjugant populations (Figure 3c). Across all three libraries, barcode frequencies in recipients were strongly correlated with their frequencies in the donor inputs, indicating that microdroplet-based conjugation does not introduce substantial skewing or selective amplification of specific variants. Although the donor libraries exhibited uneven barcode abundance distributions, as reflected by elevated Gini coefficients (Figure S5), this structure was largely preserved following conjugation, demonstrating that the platform reliably transfers complex plasmid pools even when initial variant representation is uneven. Taken together, these results show that microdroplet-based conjugation can robustly support the transfer of plasmid libraries.

### An inducible Holin/Lysin kill switch enables antibiotic-free donor counter selection

Having established that microdroplet encapsulation enhances conjugation into *S. venezuelae,* we next sought to determine whether this approach could be extended to environmental microbes and microbial communities. Initially, we sought to eliminate reliance on antibiotic-based selection. In environmental communities, introducing antibiotic resistance markers is undesirable, both for ecological and biosafety reasons and because many environmental microbes are intrinsically resistant, making antibiotic-based selection unreliable. Therefore, we first sought to remove use of antibiotics to maintain the plasmid maintenance in the donor by replacing it withauxotrophy complementation. As the donor strain carries a deletion in *dapA*, it is unable to synthesize diaminopimelic acid (DAP). By encoding the *dapA* gene on the conjugative plasmid, growth can be complemented in the absence of DAP (Figure S5). This confirms that we could effectively remove antibiotic selection for plasmid maintenance.

We next sought to remove the use of antibiotic selection for the removal of donor cells. In our prior assays, following conjugation, nalidixic acid is used to selectively remove donor cells from transconjugates. To address this, we engineered a suicide donor (SD) with a conditional holin/lysin-based kill switch circuit (Figure 4a). Following conjugation in microdroplets, donor cells are eliminated through the induction of the lysis genes that are under dual control by IPTG and AHL. The circuit also constitutively expresses mCherry and the transcriptional activator LasR, which is allosterically activated by AHL (Figure 4a). We first identified the induction conditions that maximized donor killing in bulk liquid culture. In the original study from which we obtained this circuit, the system was implemented in an *E. coli* strain that overexpresses LacI in the genome (28). Here, the donor does not carry this constitutive LacI expression cassette, so we titrated IPTG to determine whether endogenous LacI levels would affect induction. To independently track plasmid dynamics, we subsequently transformed the SD cells with a conjugative plasmid encoding GFP using the auxotrophy complementation strategy. These cells were grown overnight, subcultured into fresh media and exposed to a range of AHL and IPTG concentrations. After 3 hours of induction, cell density was measured by optical density at 600 nm (OD_600_) and fluorescence characterization of GFP was performed and normalized to cell density (FL/OD_600_) (Figure 4b, S7). From this, we identified an optimal AHL concentration of 0.5 nM that resulted in complete inhibition of growth and fluorescence, indicating effective donor killing. IPTG titration showed little to no impact under the tested conditions, but we included 1 mM IPTG in all subsequent experiments as a precaution. Because AHL is a small, diffusible molecule, its behavior in isolated microdroplets could differ from bulk conditions, potentially leading to uneven activation of the suicide circuit. Therefore, we tested the killing effectiveness of the SD cells directly within microdroplets. Cells were first grown overnight in microdroplets in the absence of AHL and IPTG. Following this initial growth, cells were decapsulated and re-encapsulated into microdroplets containing fresh media with or without AHL and IPTG and incubated overnight. SD cell survival was then assessed by microscopy, revealing robust growth in the absence of AHL and IPTG and complete loss of donor cells under the optimized inducing conditions (Figure 4c). To quantitatively validate this effect, cells were decapsulated, serially diluted, and plated for CFU enumeration, revealing an approximately four orders of magnitude reduction in viable donors upon induction (from ∼10 to ∼10³ CFU) (Figure 4d). Together, these results demonstrate efficient and antibiotic-free elimination of donor cells in microdroplets.

**Figure 4.**
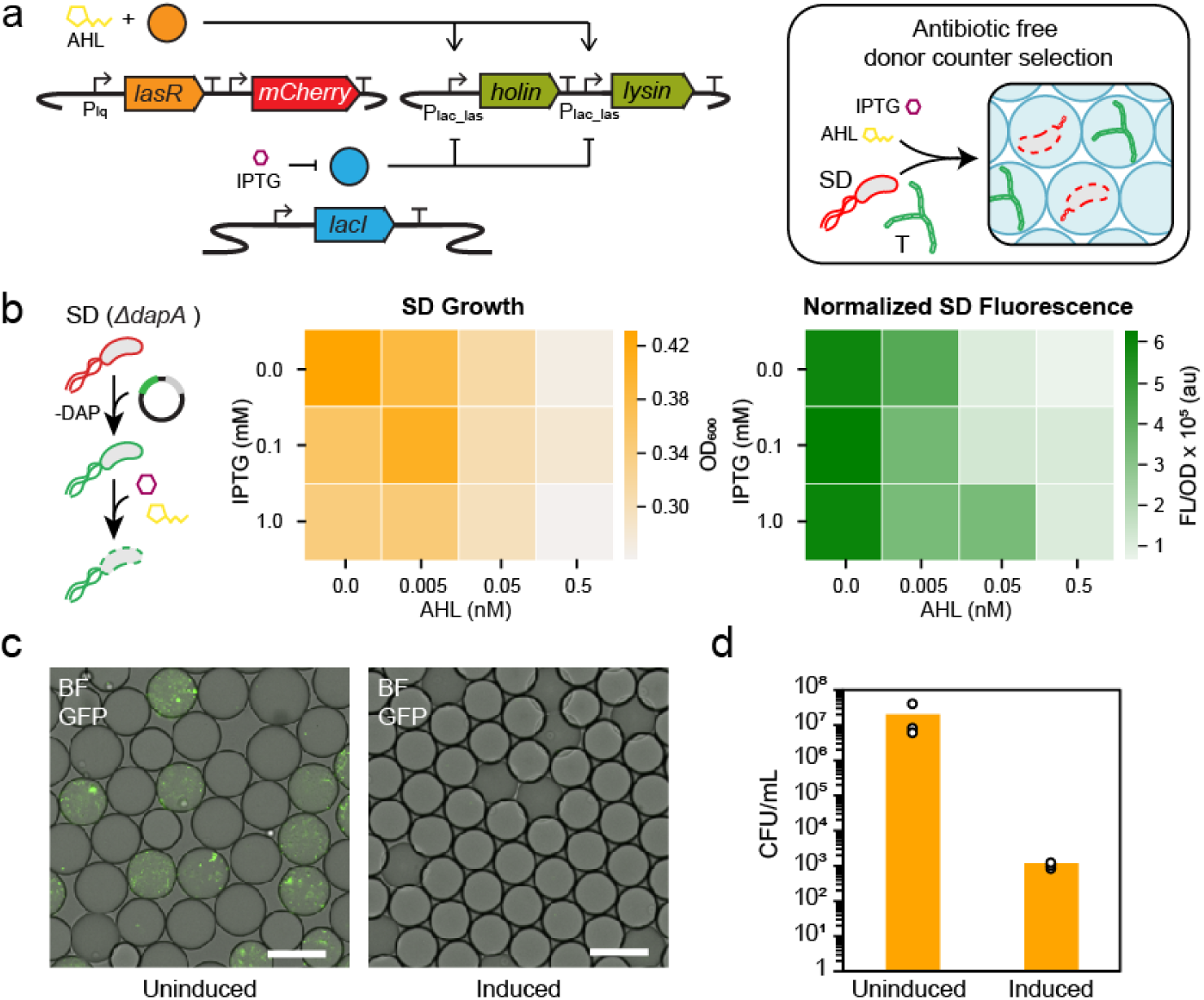
An inducible holin/lysin kill-switch for antibiotic-free donor counter selection in microdroplets. (a) Schematic of antibiotic-free donor selection via auxotrophy complementation and post-conjugation suicide donor (SD) removal via a non-conjugative, inducible holin/lysin lysis kill switch circuit triggered by AHL and IPTG. AHL activates the transcriptional regulator, LasR, and IPTG alleviates repression by endogenously encoded LacI, to induce the expression of the lysis genes, holin, and lysin. Expression of these genes leads to SD cell death. SD constitutively produces mCherry for differentiation from transconjugants that only express sfGFP from the conjugative plasmid (b) Heatmaps show the effect of AHL induction on suicide donor cell growth. Cells were grown in liquid culture overnight and subcultured into fresh media with varying concentrations of AHL. After three hours of growth, cell density was measured by optical density at 600 nm (OD_600_), and fluorescence characterization was performed (measured in units of fluorescence [FL]/optical density (OD) at 600 nm). (c) Microscope images showing suicide donor cells grown in microdroplets under induced and non-induced conditions. Cells were initially cultured and encapsulated in microdroplets, then decapsulated and re-encapsulated into fresh microdroplets at approximately one SD cell per microdroplet. Microdroplets were incubated overnight in the absence or presence of 0.5 nM AHL and 1 mM IPTG. In the absence of AHL and IPTG, microdroplets show robust donor growth, whereas AHL induction results in loss of donor cells. Scale bar: 100 µm. (d) After three days of incubation at 30°C, cells from (c) were decapsulated, serially diluted, and plated to quantify viable donor cells. In the absence of induction, donor cells remained abundant (∼10^7^ CFU), whereas induction with AHL and IPTG resulted in a ∼10^4^-fold reduction in viability (∼10^3^ CFU).

### Microdroplet-based conjugation enables DNA delivery into a soil microbial community

With the inducible suicide donor established, we next applied microdroplet-based conjugation to deliver DNA into a soil microbial community (Figure 5A). In addition to allowing for multiplexing of conjugation across diverse species, we hypothesized that by isolating individual community members into separate microdroplets and co-encapsulating them with donor cells, this strategy minimizes interspecies competition and prevents fast-growing organisms from outcompeting slower-growing or rare taxa, a common limitation of bulk liquid or plate-based methods (Dubey et al., 2021). As such we would be able to more efficiently maintain the representation of slower-growing or rare taxa and recover transconjugants that would otherwise be outcompeted or lost.

**Figure 5.**
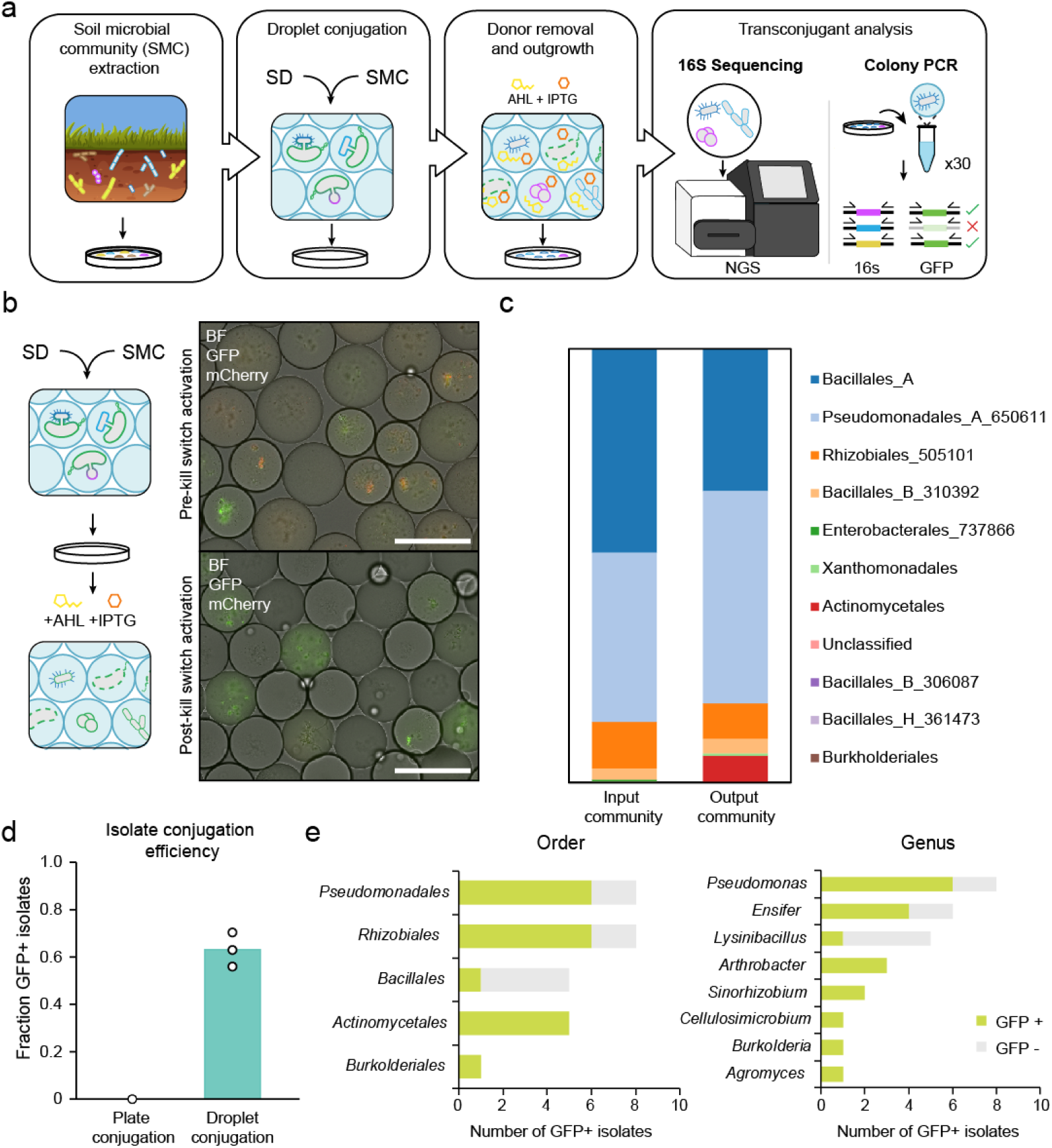
Microdroplet-based conjugation enables DNA delivery into diverse soil microbial communities. (a) Schematic of community conjugation workflow. A suicide *E. coli* donor (SD) is encapsulated with a single soil microbial community (SMC) member at a 5:1 ratio for microdroplet-based conjugation. Both donor and recipient cells are reencapsulated at an average of one cell per microdroplet with AHL and IPTG to counter select donors and enable recipient outgrowth. Following incubation, cells are serially diluted and plated for sequencing to identify plasmid uptake. (b) Green fluorescence is only observed within transconjugants following donor kill switch induction and outgrowth. (c) Community composition before and after microdroplet-based conjugation. (d) Isolate-level conjugation efficiency, calculated as the fraction of GFP-positive isolates among colonies screened by colony PCR. (e) Distribution and relative abundance of individual isolates recovered after conjugation that tested positive for the GFP sequence by colony PCR, indicating successful plasmid transfer. Each bar represents a unique isolate taxa with height corresponding to the number of GFP-positive colonies recovered for that group.

To prepare a soil microbial community, soil microbes were isolated from soil at Rice University in Houston, Texas. We selected for microorganisms compatible with the microdroplet-based conjugation workflow by growing cells in the conjugation media. Additionally, we recovered cells on media containing nalidixic acid, nystatin, and cycloheximide to inhibit fungi and favor Gram-positive bacterial species. After three days of incubation at room temperature, the microbial population was isolated from plates by washing with 1 mL of the conjugation media and glycerol stocked to allow for use across multiple experimental replicates.

To assess microdoplet-based conjugations, the suicide donor was co-encapsulated with this soil microbial community at a 5:1 donor-to-recipient ratio to ensure that virtually every microdroplet contained at least one donor cell. After an initial conjugation period, cells were re-encapsulated at approximately one recipient cell per microdroplet in the presence of AHL and IPTG to trigger the holin/lysin kill switch, removing donors and allowing recipient cells to grow. Successful DNA transfer was observed via GFP expression in recipient cells. Prior to induction of the holin/lysin kill switch, microdroplets displayed both GFP and mCherry fluorescence, reflecting the presence of donor cells alongside transferred plasmids. Following AHL and IPTG induction, donor cells were effectively eliminated, and fluorescence was restricted to GFP only in recipient cells (Figure 5B).

After incubation, cells were serially diluted and plated for further analysis. We first aimed to understand if community composition changed before and after microdroplet-based conjugation. To do this, total DNA was extracted from the original community and the post conjugation community and 16S rRNA sequencing was performed. From this we saw that community composition was largely unchanged following conjugation in microdroplets. Eleven unique bacterial orders were detected in both the original and post-conjugation samples. The dominant populations were *Pseudomonadales* and *Bacillales A*, comprising approximately 30-50% of the community, while *Actinomycetales* accounted for approximately 0.1-6%. Notably, *Actinomycetales* include slow-growing taxa such as *Streptomyces* indicating that these organisms were retained throughout the conjugation process. This suggests that microdroplet-based conjugation does not strongly bias against slower-growing or less competitive community members, preserving taxa that are often underrepresented in bulk mating experiments.

To understand the overall conjugation efficiency into the soil microbial community and identify which community members were conjugated, we next performed colony PCR on 30 isolates to quantify the fraction of isolates containing the GFP-encoding plasmid. Surprisingly, we saw that ∼60% of the total isolates were GFP positive, suggesting a surprisingly high conjugation efficiency using our method (Figure 5d, S7). To ascertain the identity of these transconjugates we performed 16S ribosomal RNA (rRNA) sequencing on the GFP-positive and -negative isolates. Transconjugates spanned five bacterial taxonomic orders, including *Actinomycetales, Bacillales, and Pseudomonadales* (Figure 5e).

At the genus level, the 30 recovered isolates represented eight distinct taxa. Many of these genera are common in soil and have well-documented ecological roles and biotechnological potential. *Pseudomonas* and *Burkolderia* species have been explored extensively in biodegradation and metabolic engineering, including heterologous production of complex metabolites (29). In contrast, *Ensifer* and *Sinorhizobium* are emerging chassis, traditionally studied as nitrogen-fixing symbionts; they are only beginning to be explored for genetic manipulation beyond symbiotic traits (30). Finally, genera such as *Arthrobacter,* known for their resistance to harsh climates and heavy metal sequestration, remain largely untapped for synthetic biology due to low tractability and limited toolsets (31). These results demonstrate that microdroplet-based conjugation can access a phylogenetically and functionally diverse set of organisms, including taxa that are typically difficult to manipulate using conventional approaches.

As a control, the same community was incubated with donor cells using conventional plate-based conjugation followed by bulk liquid outgrowth. In this no-microdroplet condition, none of the thirty randomly screened isolates tested positive for GFP (Figure 5d), and only one taxonomic order, *Bacillales,* made up a majority of the isolates (Figure S8). These results highlight that microdroplet spatial segregation is essential both for successful plasmid transfer and for preserving slower-growing taxa that are lost in bulk conjugation approaches.

## Discussion

A central bottleneck in microbial engineering is the efficient delivery of DNA into diverse and poorly characterized non-model hosts. This is particularly true in cases where the introduction of large DNA libraries is needed, for example, pooled CRISPR screens (32, 33). Transformation and conjugation efficiencies vary widely across species and are difficult to predict (34), limiting DNA delivery as a generalizable engineering step. This challenge is compounded by two constraints, the inefficiency to construct strain libraries, and the identification of genetically tractable hosts, which remains largely dependent on low-throughput, trial and error screening of individual isolates, or expensive automated workflows (35). Droplet microfluidics address these limitations by compartmentalizing cells into picoliter-scale microdroplets that enforce and stabilize microbial co-localization for conjugation while preserving high experimental throughput. In this format, conjugation is transformed from a bulk population process into a massively parallel set of isolated microenvironments, enabling both efficient DNA delivery and simultaneous interrogation of diverse host-vector combinations.

The microfluidic technology of this platform supports seamless integration with complementary droplet-based techniques across the workflow. Upstream, controlled droplet merging provides a mechanism to precisely regulate donor-recipient interactions through temporal control over the initiation of conjugation (36). While downstream, fluorescence-activated droplet sorting (FADS) would allow enrichment of rare successful transfer events directly within droplets, by coupling plasmid transfer to the expression of a fluorescent reporter in transconjugates (37). When combined with a synthetic donor removal circuit, this would establish an antibiotic-free enrichment strategy, where fluorescence provides a positive selection for transconjugates, while donor depletion suppresses background signal. This combined approach replaces reliance on host-specific antibiotic markers or engineered auxotrophies with a more generalizable, phenotype-based selection strategy. Further, droplet dispensing technology enables the recovery and isolation of individual transconjugates for downstream characterization (38). Together, these capabilities would establish a continuous, scalable pipeline from DNA delivery to selection and strain isolation at scale.

We demonstrate that microdroplet spatial confinement provides enhanced conjugation efficiency, which we propose is by stabilizing the donor-recipient interactions required for successful plasmid transfer. This is particularly impactful for engineering of individual non-model microorganisms, where low baseline transfer efficiency often prevents genetic manipulation altogether. By increasing DNA delivery within individual species, this platform expands the access to organisms that are otherwise considered genetically intractable and enables workflows that require high numbers of independent transfer events, including plasmid library approaches. Library bottlenecking remains a major limitation in pooled functional genomics approaches such as CRISPR interference (CRISPRi) and activation (CRISPRa) perturbation screens, where large libraries are required to uncover gene-phenotype relationships at the genome scale (39). CRISPRi and CRISPRa systems have recently been demonstrated in *S. venezuelae* for targeted gene modulation and interrogation of antimicrobial biosynthetic pathways (40), highlighting their potential for functional genomics in Actinomycetes. However, scaling these approaches to genome-wide perturbations across diverse hosts remains constrained by inefficient DNA delivery. In practice, achieving sufficient library representation has often required multiple conjugation steps or parallelized mating workflows in multiwell formats (Clarke et al., 2026; Li et al., 2026), reflecting the limited scalability of conventional DNA transfer methods. By enabling high-efficiency delivery of plasmid libraries in a highly parallelized format, this platform provides a scalable route toward genome-wide functional genomics in non-model organisms.

Beyond individual microbes, we anticipate this platform will be of value for studies and engineering of microbial communities. For example, while DNA can be directly transferred into complex microbial communities (14, 43), performing these experiments in bulk populations is complicated by growth rate disparities and interspecies competition, which bias toward fast-growing taxa, limiting the range of accessible recipients. Droplet compartmentalization mitigates these effects by isolating the recipient population into a large set of parallel conjugation reactions. This reduces competition during DNA transfer, enabling a broader sampling of recipient diversity. While this study focused on a selectively enriched soil community, this approach is readily extended to more complex environments, where increased diversity may further probe conjugation permissivity.

We envision that this platform can be extended as a modular system in which donor identity, plasmid architecture, and community composition can be easily varied. This flexibility enables high efficiency DNA delivery across diverse microbial hosts. The platform can be used to interrogate horizontal gene transfer dynamics, such as quantifying plasmid host range, and in future interactions can be coupled with emerging molecular recording strategies that enable high-resolution tracking of gene transfer events in complex communities (14). Collectively, this works established microdroplet-based conjugation as a programmable strategy that transforms DNA delivery from a limiting step into a controllable experimental variable, enabling access to DNA transfer across both individual hosts and complex communities.

## Materials and Methods

### Bacterial strains and growth conditions

*E. coli* strains were grown in Luria Bertani (LB) broth, or LB agar (1.5%). 2,6-diaminopimelic acid (DAP) was added to LB at a final concentration of 0.1 mM for culturing strains derived from *E. coli* WM6029. *S. venezuelae* ATCC 10712 were cultured in complete supplement medium (CSM) or maltose, yeast extract, malt extract (MYM) agar. To make CSM, 30 g of tryptic soy broth, 1.2 g of yeast extract and 1 g of MgSO4 were dissolved in 1 L of water. During use, 1% (v/v) of filter-sterilized 50% glucose and 40% maltose were supplemented. MYM media was prepared using 2.0 g Maltose, 2.0 g yeast extract, and 5.0 g Malt extract in 500 ml water. All conjugations were conducted using AS-1 medium or agar. AS-1 base was prepared by adding 1.25 g of soluble starch, 0.625 g of NaCl, 0.25 g of yeast extract, to 250 ml of water. After autoclaving, the AS1 base was mixed with 25 ml of autoclaved 10% (w/v) Na2SO4 and 5 ml of a filter sterilized ARN amino acid solution (1 g of alanine, 1 g of arginine, and 1 g or asparagine dissolved in 50 ml sterile water). When necessary, media was supplemented with antibiotics (50 ug/ml apramycin, 100 ug/ml carbenicillin, 100 ug/ml kanamycin, nalidixic acid 30 ug/ml).

### Plasmid and library construction

All plasmids and primers can be found in Table S1 and Table S2. All plasmids were constructed with either Gibson assembly (Gibson et al., 2009), Golden Gate assembly (Engler et al., 2008) or PCR. Plasmid construction was performed in *E. coli* strain NEB Turbo unless otherwise noted. The barcoded plasmid library was generated using an oligonucleotide containing a 10-nucleotide degenerate barcode (N_10_) flanked by BbsI recognition sites, with a primer binding site on one end. The oligonucleotide was subjected to a single-cycle PCR extension using a complementary primer to generate double-stranded DNA. The resulting product was purified and used as an insert in a Golden Gate assembly reaction to introduce the barcode library into the conjugative plasmid backbone. The assembled library was transformed directly into *E. coli* WM6029 for downstream conjugation experiments.

### Fabrication of microfluidic device

Microfluidic devices were fabricated using standard soft lithography techniques (Xia & Whitesides, 1998). SU-8 photoresist was patterned on silicon wafers to generate channel molds. The microfluidic devices were made using PDMS (poly(dimethylsiloxane)) at a 10:1 (w/w) base to curing agent ratio. The PDMS was degassed and cured at 80 °C for 2-18 hours. The cured PDMS was peeled off the mold and inlet holes for tubing were punched using 0.5 mM core sampling puncher. The PDMS devices were bonded to glass slides following oxygen plasma treatment and the channels were made hydrophobic by incubating the devices at 80C for two days.

### Cell encapsulation

The microfluidic microdroplet generator has an inlet for the oil phase and two inlets for the two different bacterial cultures that are targeted to the flow-focusing junction to form uniform microdroplets. Aqueous flow rates of 15 μL/min and an oil flow rate of 100 μL/min was used to produce microdroplets of 90 μm in diameter. Bacterial cultures were used as the aqueous phase, whereas Novec HFE 7500 fluorinated oil (3M, Maplewood, MN) with either 1.5% PicoSurf surfactant (Sphere Fluidics, Cambridge, U.K.) or 1.75% Microdroplet imaging was done using a VitroCom glass capillary tube (Mountain Lakes) under a 100x objective on a Ti2E inverted microscope (Nikon). Fluorescence was imaged using either GFP: Excitation: 480/30 nm, Emission: 535/20 nm or mCherry: Excitation: 560/40 nm, Emission: 635/60 nm filters (Chroma).

### Benchtop and Microfluidic Conjugations

Donor strains harboring conjugative plasmids were cultured in LB supplemented with selective antibiotics or DAP as required. Overnight cultures were grown at 37 °C with shaking (225 rpm). Cells were harvested at stationary phase (OD600 ≈ 1.0) and were washed and concentrated 10-fold using fresh medium to remove antibiotics. Cells were then diluted at defined cell densities with ratios chosen to achieve desired occupancy, assuming Poisson loading assumptions for microdroplet experiments and applied equivalently in benchtop conjugations. Benchtop conjugations were performed by mixing donor and recipient cells at equal volumes. For S. venezuelae conjugations, an equal volume of thawed spore glycerol stocks were used as the recipient. Mixtures were plated to either AS1 agar supplemented with 0.1 mM DAP and incubated at 30°C 24 h. Microdroplet-based conjugation experiments were conducted using flow-focusing geometries to generate aqueous microdroplets in an oil phase containing 1.75% (w/w) biocompatible surfactant. Donor and recipient cells were co-encapsulated within microdroplets at the same input ratios. Flow rates were controlled using syringe pumps to produce microdroplets with diameters of approximately 90 μm. Microdroplets were collected and incubated at 30 °C for defined time periods to allow conjugation to occur.

Following incubation, emulsions were broken using a detergent (1H, 1H, 2H, 2H-Perfluoro-1-octanol), and 100 µl of cells were transferred to AS1 agar for 24 hours. For all conjugations, following incubation on AS1 agar, cultures were harvested, serially diluted and plated on selective AS1 agar to quantify transconjugants, and recipients. Conjugation efficiency was calculated as the ratio of transconjugants to total recipient cells (CFU).

### Donor growth measurements

*E. coli* colonies were grown in 5ml LB supplemented with kanamycin or ampicillin as needed at 37 C with shaking (225 rpm) overnight. Cultures were then diluted 1:50 into fresh LB containing the appropriate antibiotics and transferred to either a 96-well microplate (150 µl per well) or a deep-well block (300 µl per well).

For growth curves experiments, cultures were then grown in a microplate reader (Tecan Spark multimode plate reader) at 37 C with orbital shaking (90 rpm) for 24 h, and OD_600_ measurements were recorded every 10 min. For endpoint measurements, cultures were grown in a Vortemp shaker at 37 C for 3 hours following subculturing. Subsequently, 100 µl of each culture was transferred to a 96-well plate, and OD_600_ was measured using a microplate plate reader. For data analysis, OD_600_ values of the media-only controls were averaged and subtracted from corresponding measurements for each condition.

### Barcoded DNA Library Analysis

Raw FASTQ files were processed using a custom Python-based pipeline to extract and quantify barcode sequences from donor and recipient populations. Briefly, sequencing reads were scanned for the presence of a fixed anchor sequence (GGGCCAGGGCCGTTG), allowing up to one mismatch to account for sequencing errors. Upon identification of this sequence, a 10-nucleotide barcode immediately downstream was extracted. Barcodes were filtered to remove low-confidence observations. Donor barcodes were required to have a minimum of two supporting reads, while recipient barcodes required at least one read, that was also detected in the donor population.

### Soil community Preparation

Soil samples were collected from the surface layer (≤2 cm depth) of a site located in Rice University, Houston, TX. Approximately 5 g of soil was suspended in 45 mL of sterile 0.9% (w/v) NaCl solution, prepared by dissolving 0.45 g NaCl in 50 mL sterile water and autoclave sterilized. The suspension was gently vortexed at room temperature for 1 h to dislodge microbial cells, followed by incubation at room temperature for 10 min to allow large particulates to settle. The resulting supernatant was collected and serially diluted 10-fold and of each dilution (10^0^-10^−3^) were spread onto AS1 agar plates supplemented with nalidixic acid (30 µg/mL), cycloheximide (50 µg/mL), and nystatin solution to suppress fungal growth and enrich for gram positive bacterial populations. Plates were ncubated at room temperature for 3 days. Following incubation, microbial biomass was recovered by washing plate surfaces with 1 mL sterile phosphate-buffered saline (PBS). Suspended cells from replicate plates were pooled and mixed with sterile 50% (v/v) glycerol at a 1:1 ratio to generate glycerol stocks.

### Soil Community Amplicon and Colony PCR Sequencing for 16S rRNA and GFP

Microbial community composition was assessed by 16S rRNA gene amplicon sequencing. Following incubation, cells were harvested from agar plates by washing the surface with 1 mL sterile PBS. The resulting suspensions were collected, and genomic DNA was extracted using a commercial DNA extraction kit according to the manufacturer’s instructions. Amplicon libraries were generated by PCR amplification of the 16S rRNA gene using primers containing Illumina adapter sequences. PCR products were purified and submitted for sequencing using the Amplicon-EZ service (Genewiz, Azenta Life Sciences). To assess DNA transfer and isolate identity, individual colonies obtained following conjugation were screened by colony PCR. Colonies were picked and resuspended in 40 µL sterile water, and 5 µL of the suspension was used as template in PCR reactions targeting either the GFP gene or the 16S rRNA gene. PCR products were analyzed by agarose gel electrophoresis. Representative amplicons were purified and submitted for Sanger sequencing to confirm gene presence (GFP) or determine taxonomic identity (16S).

### 16S analysis

Raw paired-end sequencing reads were processed using Cutadapt to remove primer sequences and low-quality regions. Primer trimming was performed using the forward primer sequence (TACTAACGCGAAGAACCTTAC) and reverse primer sequence (GACGGGCGGTGWGTRCA), corresponding to the 16S rRNA gene amplification region. Primer matching required a minimum overlap of 5 nucleotides, and reads were trimmed using anchored 5′ adapter matching. Reads that did not contain recognizable primer sequences were written to separate “untrimmed” files. Additionally, a minimum length threshold of 400 bp was enforced following trimming to remove short reads. Trimmed sequences were imported into QIIME 2 using a paired-end manifest format. Sequence quality was assessed using the demultiplexing summary visualization, which indicated high-quality reads and did not require additional truncation. Denoising, dereplication, chimera removal, and feature table construction were performed using the DADA2 plugin with paired-end processing. Parameters included no truncation of forward or reverse reads, a maximum expected error threshold of 4 for both directions, a minimum overlap of 15 bp for read merging, and pseudo-pooling to improve sensitivity to rare variants. This process generated a feature table and representative sequences corresponding to amplicon sequence variants (ASVs). Feature tables, representative sequences, and denoising statistics were visualized using QIIME 2 visualization tools. A phylogenetic tree was constructed from the representative sequences using MAFFT alignment followed by FastTree, implemented through the QIIME 2 phylogeny pipeline. Diversity analyses were performed using the core-metrics-phylogenetic workflow at a sampling depth of 94,935 sequences per sample, based on the minimum observed sequencing depth. Alpha rarefaction analysis was also conducted with a maximum depth of 4,000 sequences. To further refine the dataset, representative sequences were exported and filtered to remove ASVs shorter than 400 bp. Taxonomic classification was performed using a naïve Bayes classifier trained on a full-length 16S rRNA reference database (2024.09 backbone release), implemented via the QIIME 2 feature-classifier plugin. Taxonomic assignments were visualized using QIIME 2 metadata tools.

## Author information

### Author contributions

A.S, J.C, and Y.S conceived the study. A.S. and A.T. performed all experiments and data analysis. A.S., J.C., and Y.S. wrote the manuscript. All authors reviewed the manuscript.

## Notes

The authors declare no competing financial interest.

## Acknowledgements

This project was supported by the National Science Foundation Award MCB-2247573 (J.C., Y.S.).

## Supplemental information

**Supplementary Table 1.**
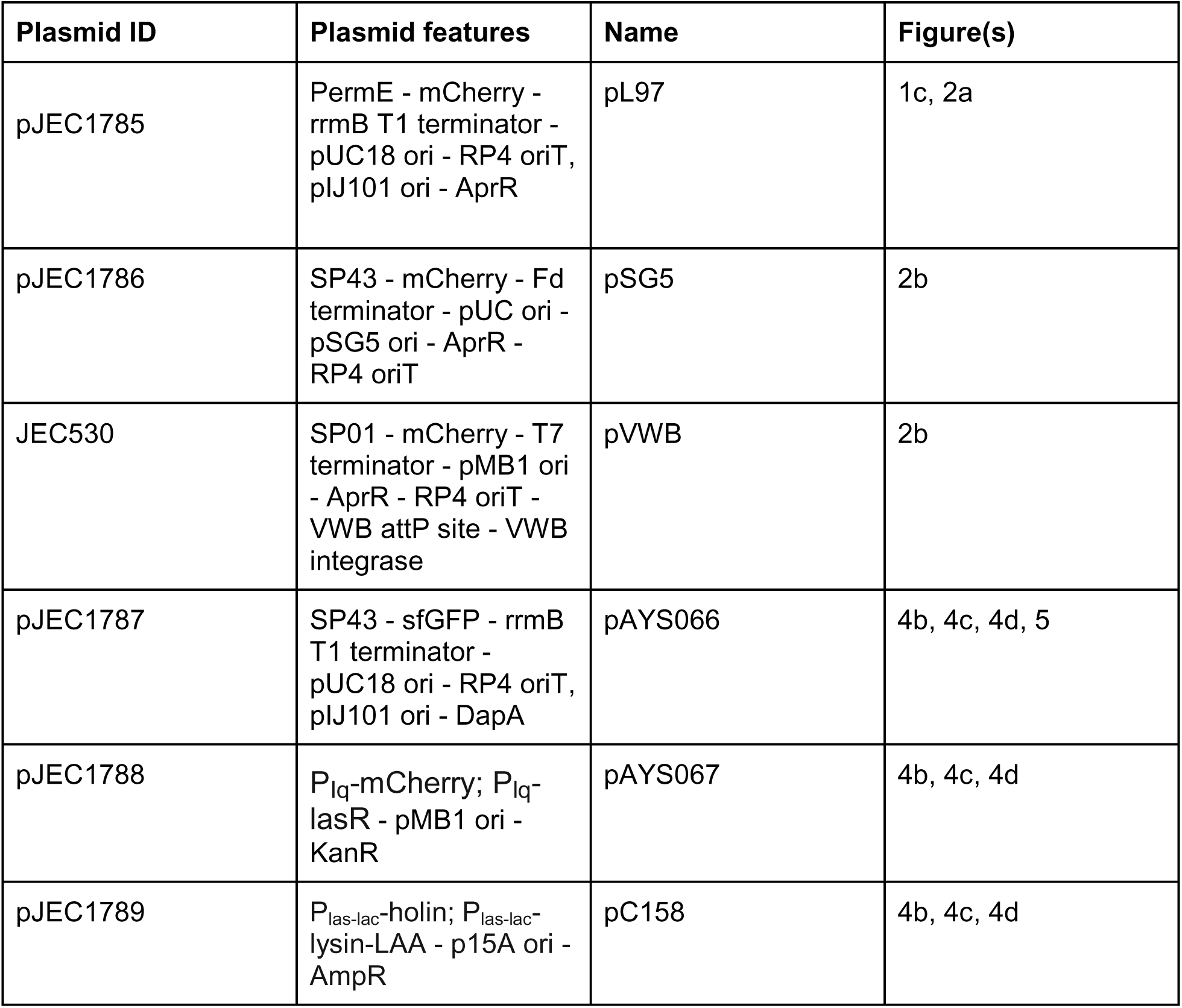
List of all plasmids used in this study.

**Supplementary Table 2.**
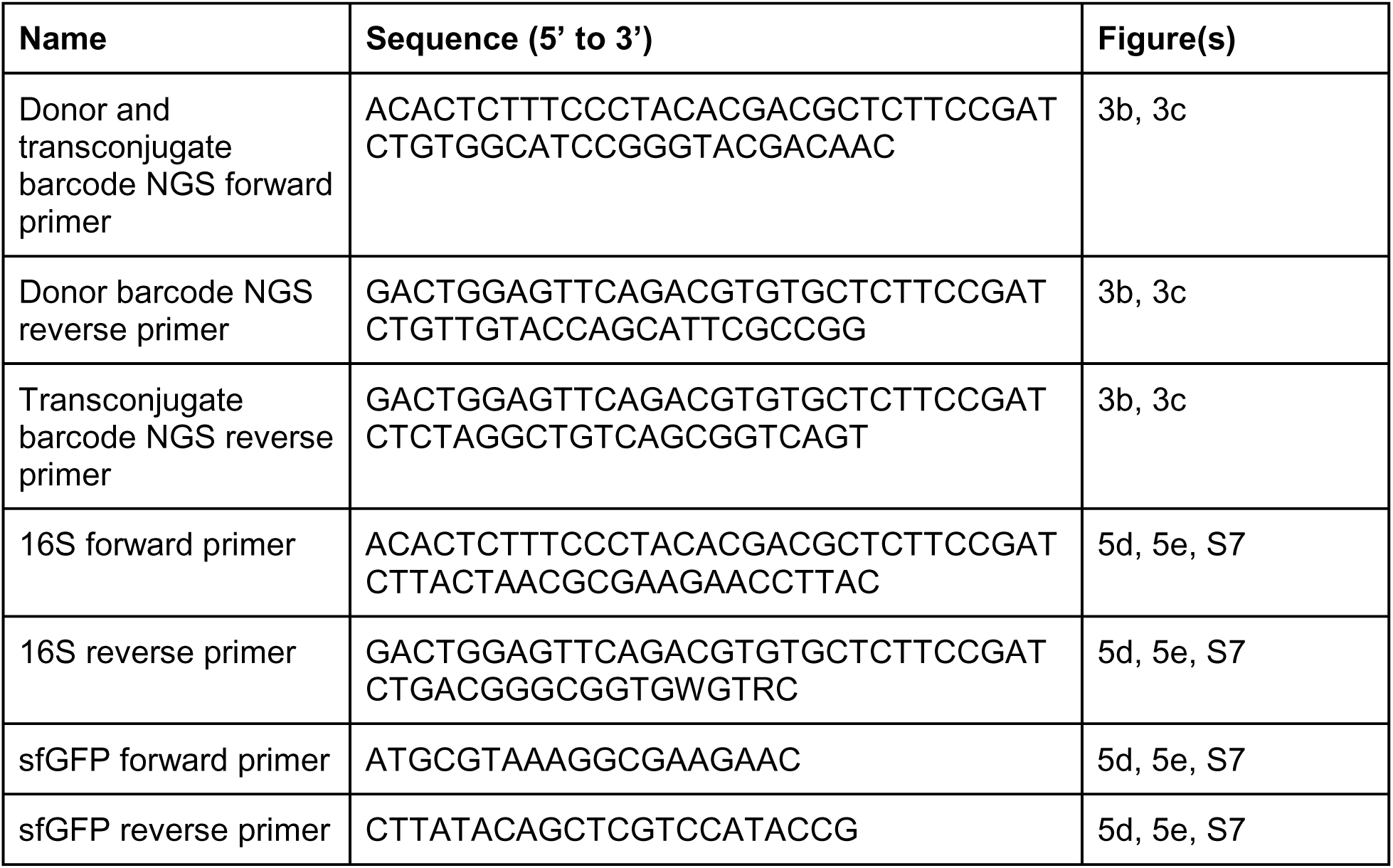
List of relevant Oligos used in this study.

**Figure S1.**
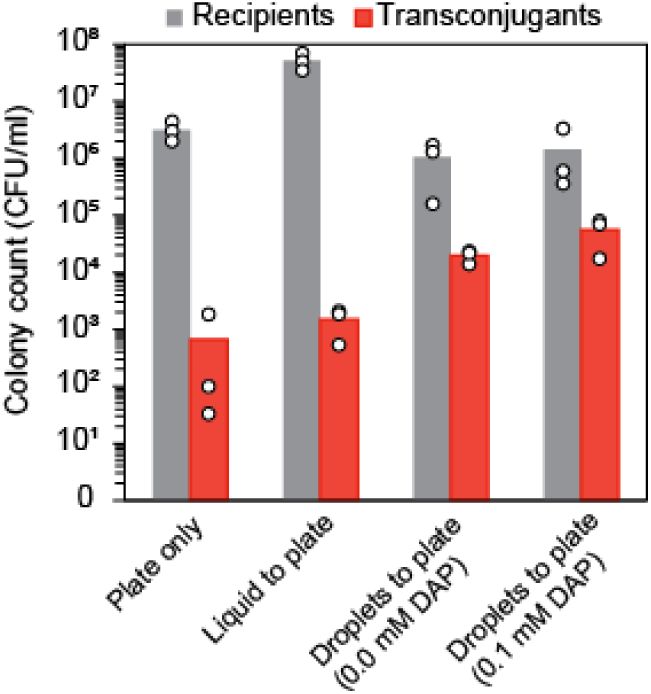
Recipient and transconjugant colony-forming units (CFU) used to calculate conjugation efficiency values shown in Figure 1. CFU were quantified by plating onto selective media for recipients and transconjugants, respectively. *S.venezuelae* recipients were selected by plating onto AS1 media supplemented with nalidixic acid to inhibit donor growth, while transconjugants were selected by the addition of apramycin to the media. Plates were incubated at 30 °C for 2-5 days and the resulting colonies were counted. Compared to cells incubated on plates, incubation in microdroplets produced between 10 and 100-fold increase in the absolute number of transconjugates.

**Figure S2.**
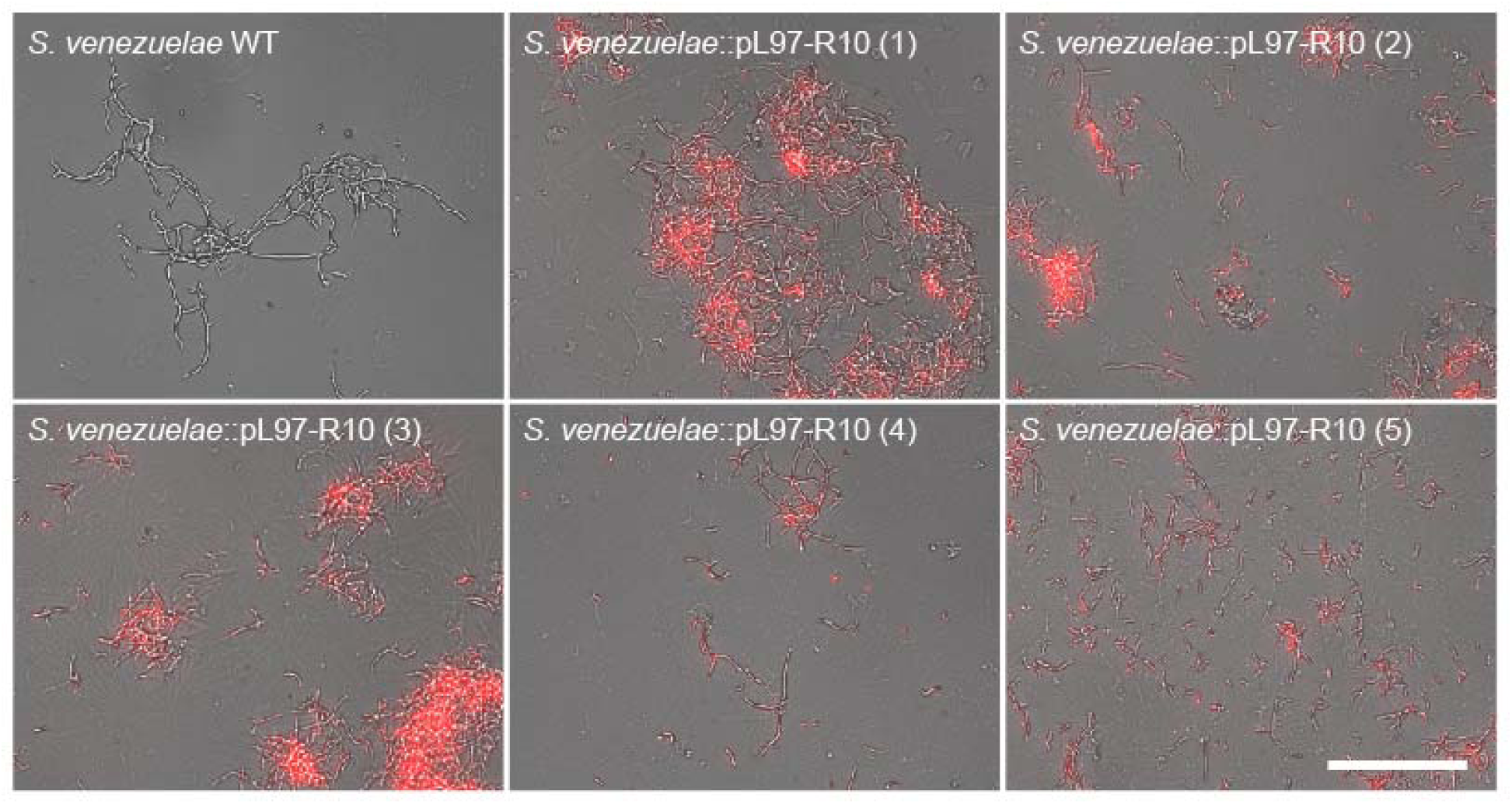
Fluorescence microscopy confirms mCherry expression in *S. venezuelae* transconjugants. Representative fluorescence microscopy image of wild-type *Streptomyces venezuelae* (WT) and five independently isolated transconjugant colonies. Following microdroplet-based conjugation transconjugant *S.venezuelae* colonies were imaged for mCherry fluorescence. Wild-type cells show no detectable red fluorescence, while transconjugants exhibit mCherry fluorescence, confirming successful plasmid transfer and expression. Scale bars, 100 µm

**Figure S3.**
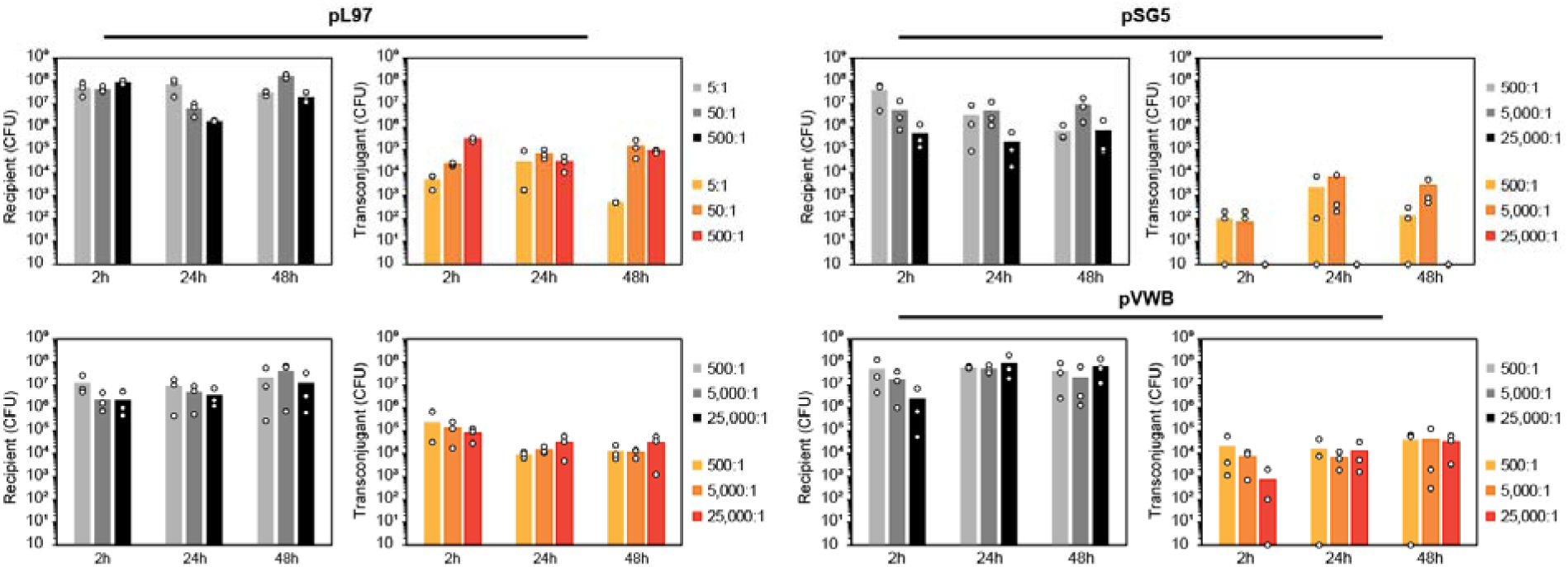
Recipient and transconjugant colony-forming units (CFU) used to calculate conjugation efficiency values shown in Figure 2. Recipient and transconjugant colony-forming units (CFU) were measured across varying donor to recipient ratios and incubation times in microdroplets. For pL97, donor-to-recipient ratios of 5:1, 50:1, 500:1, 5,000:1, and 25,000:1 were tested, while pSG5 and pVWB were only evaluated at ratios of 500:1, 5,000:1, and 25,000:1. Cells were incubated in microdroplets for 2 h, 24 h, or 48 h prior to plating. CFU were quantified by selective plating, with recipients selected on AS1 media supplemented with nalidixic acid and transconjugants selected with nalidixic acid and apramycin. Plates were incubated at 30 °C for 2–5 days before colony counting.

**Figure S4.**
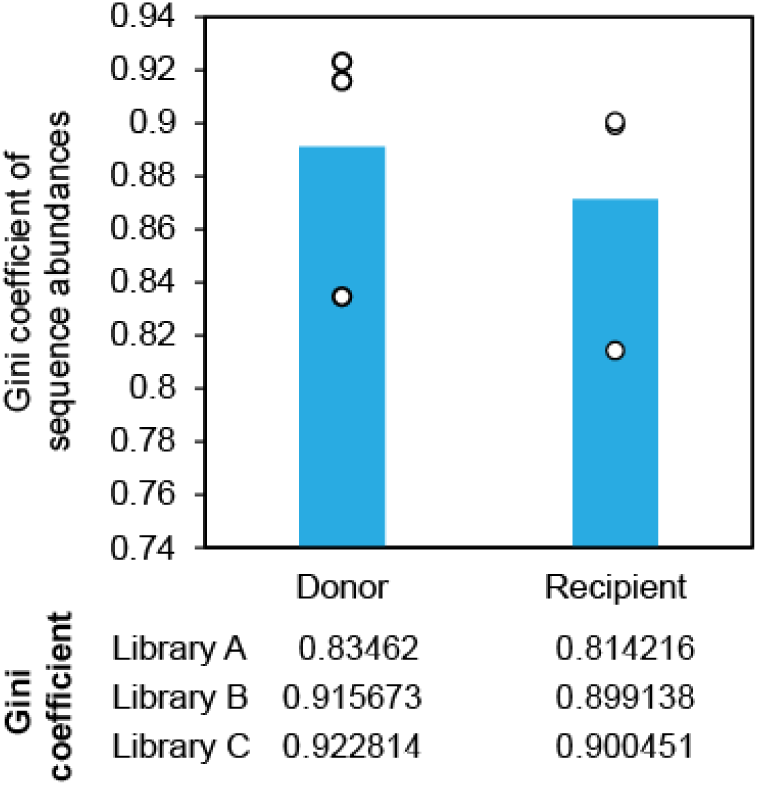
Gini coefficients of barcoded plasmid libraries. Comparison of barcode library uniformity before and after conjugation, quantified using the Gini coefficient of sequence abundances. Input libraries were sequenced from donor minipreps (Donor), and corresponding output libraries were recovered by PCR amplification from transconjugant *S. venezuelae* genome following microdroplet-based conjugation (Recipient). Each point represents an independently cloned library.

**Figure S5.**
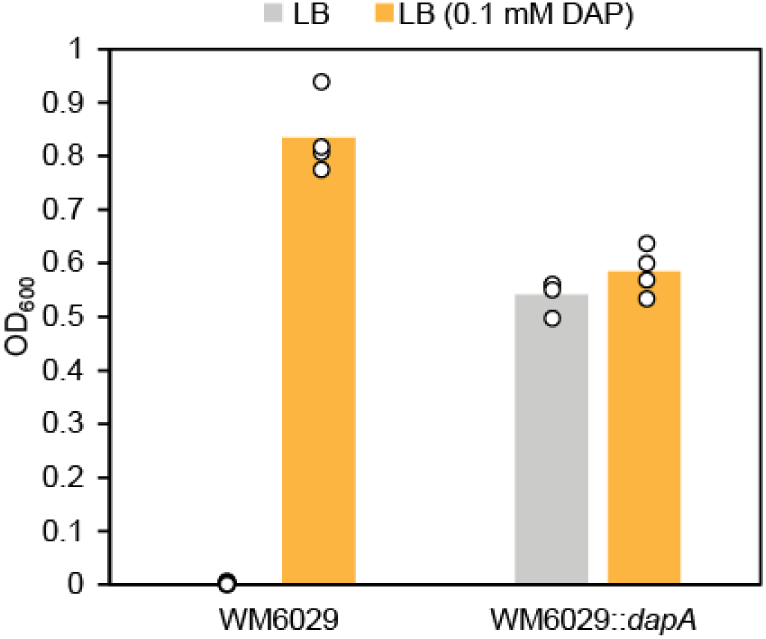
Evaluating *dapA* complementation on donor growth. *E. coli* WM6029 were transformed with a plasmid that either contained or lacked *dapA*. Following an overnight incubation at 37 °C, optical densit at 600 nm (OD_600_) was measured. Cells that are not complemented with *dapA* are only able to grow in the presence of 0.1mM DAP, while in the absence of DAP, there was no difference in growth for the *dapA* complemented strain.

**Figure S6.**
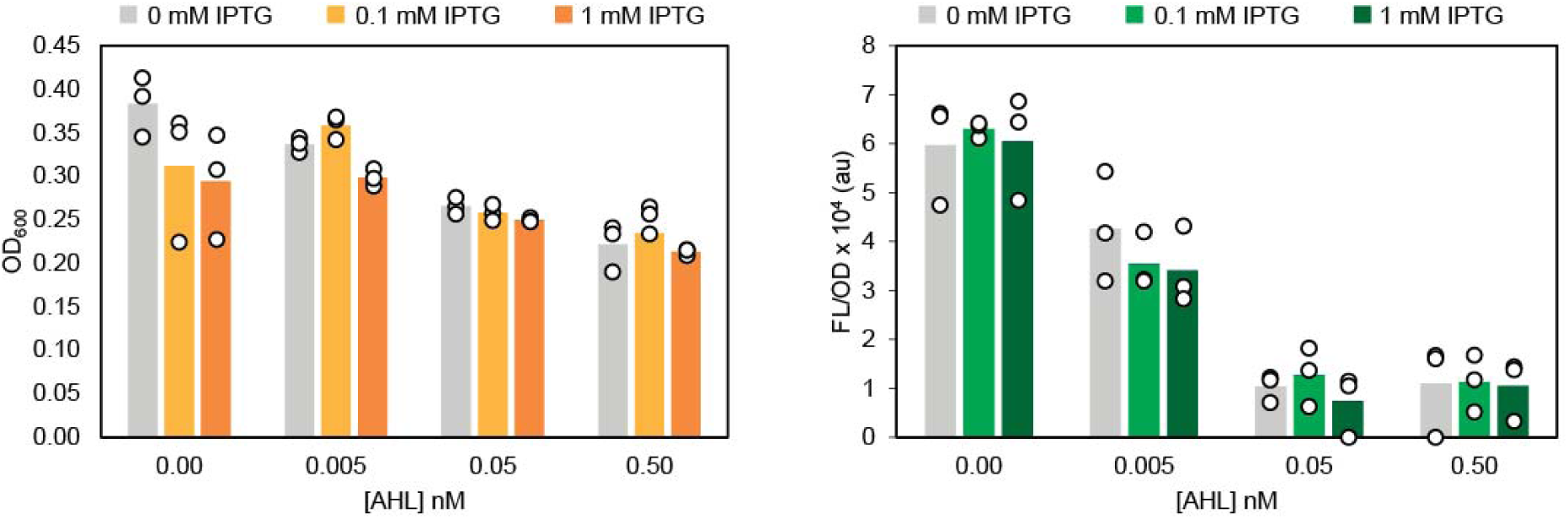
Suicide donor AHL/IPTG titration FL/OD raw values. To initially characterize AHL and IPTG induction for efficient donor removal via a holin and lysin mechanism, *E. coli* WM6026 was transformed with three plasmids: the two kill switch circuit plasmids and a conjugative plasmid that expresses dapA and sfGFP. Cells were grown in liquid culture overnight and subcultured into fresh media with varying concentrations of AHL (0 nM - 0.5 nM) and IPTG (0 mM - 1 mM). Cell density was measured after three hours of growth by optical density at 600 nm (OD600), and fluorescence characterization of the conjugative plasmid was performed (measured in units of fluorescence/optical density (FLOD).

**Figure S7.**
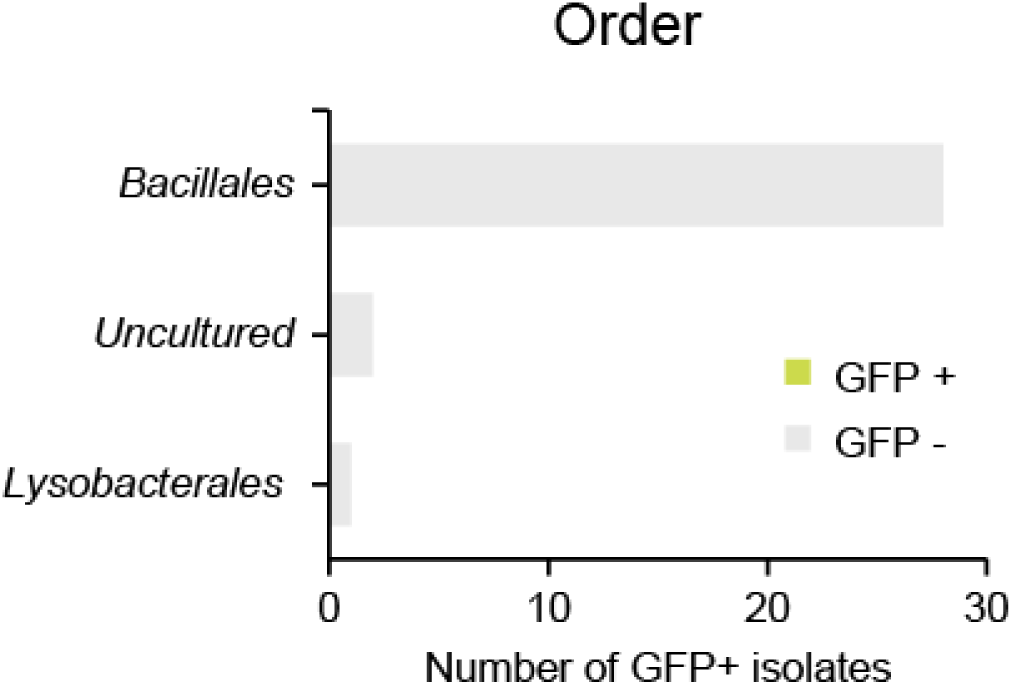
Colony PCR analysis of isolates from bulk community conjugation control. To assess plasmid transfer under conventional plate-based conjugation conditions and bulk liquid culture outgrowth, the soil microbial community was incubated with suicide donor cells on solid media without microdroplet encapsulation. After conjugation, cells were transferred to liquid media with AHL and IPTG to eliminate donor cells and allow transconjugant outgrowth. Following this growth period, cultures were serially diluted and plated, and 30 randomly selected isolates were screened by colony PCR for the presence of the GFP on the conjugative plasmid. No isolates tested positive for GFP, indicating a lack of detectable plasmid transfer. 16S sequencing of the isolates revealed that *Bacillales* were the single taxonomic order that was predominantly recovered, highlighting that bulk conjugation is both less efficient and less capable of preserving slower-growing or less competitive taxa.

