## Supplementary Material for "A droplet microfluidic-based platform for enhanced DNA delivery in non-model organisms"

**TABLE OF CONTENTS**

| **Page 2** | Supplementary Table 1. List of all plasmids used in this study. |
| --- | --- |
| **Page 3** | Supplementary Table 2. List of relevant oligos used in this study. |
| **Page 4** | Figure S1. Recipient and transconjugant colony-forming units (CFU) used to calculate conjugation efficiency values shown in Figure 1. |
| **Page 5** | Figure S2. Fluorescence microscopy confirms mCherry expression in *S. venezuelae* transconjugants. |
| **Page 6** | Figure S3. Recipient and transconjugant colony-forming units (CFU) used to calculate conjugation efficiency values shown in Figure 2. |
| **Page 7** | Figure S4. Gini coefficients of barcoded plasmid libraries. |
| **Page 8** | Figure S5. Evaluating *dapA* complementation on donor growth. |
| **Page 9** | Figure S6. Suicide donor AHL/IPTG titration FL/OD raw values. |
| **Page 10** | Figure S7. Colony PCR analysis of isolates from bulk community conjugation control. |

Supplementary Table 1. List of all plasmids used in this study.

| **Plasmid ID** | **Plasmid features** | **Name** | **Figure(s)** |
| --- | --- | --- | --- |
| pJEC1785 | PermE - mCherry - rrmB T1 terminator - pUC18 ori - RP4 oriT, pIJ101 ori - AprR | pL97 | 1c, 2a |
| pJEC1786 | SP43 - mCherry - Fd terminator - pUC ori - pSG5 ori - AprR - RP4 oriT | pSG5 | 2b |
| JEC530 | SP01 - mCherry - T7 terminator - pMB1 ori - AprR - RP4 oriT - VWB attP site - VWB integrase | pVWB | 2b |
| pJEC1787 | SP43 - sfGFP - rrmB T1 terminator - pUC18 ori - RP4 oriT, pIJ101 ori - DapA | pAYS066 | 4b, 4c, 4d, 5 |
| pJEC1788 | P_Iq_-mCherry; P_lq_-lasR - pMB1 ori - KanR | pAYS067 | 4b, 4c, 4d |
| pJEC1789 | Plas-lac-holin; Plas-lac-lysin-LAA - p15A ori - AmpR | pC158 | 4b, 4c, 4d |

Supplementary Table 2. List of relevant Oligos used in this study.

| **Name** | **Sequence (5’ to 3’)** | **Figure(s)** |
| --- | --- | --- |
| Donor and transconjugate barcode NGS forward primer | ACACTCTTTCCCTACACGACGCTCTTCCGATCTGTGGCATCCGGGTACGACAAC | 3b, 3c |
| Donor barcode NGS reverse primer | GACTGGAGTTCAGACGTGTGCTCTTCCGATCTGTTGTACCAGCATTCGCCGG | 3b, 3c |
| Transconjugate barcode NGS reverse primer | GACTGGAGTTCAGACGTGTGCTCTTCCGATCTCTAGGCTGTCAGCGGTCAGT | 3b, 3c |
| 16S forward primer | ACACTCTTTCCCTACACGACGCTCTTCCGATCTTACTAACGCGAAGAACCTTAC | 5d, 5e, S7 |
| 16S reverse primer | GACTGGAGTTCAGACGTGTGCTCTTCCGATCTGACGGGCGGTGWGTRC | 5d, 5e, S7 |
| sfGFP forward primer | ATGCGTAAAGGCGAAGAAC | 5d, 5e, S7 |
| sfGFP reverse primer | CTTATACAGCTCGTCCATACCG | 5d, 5e, S7 |

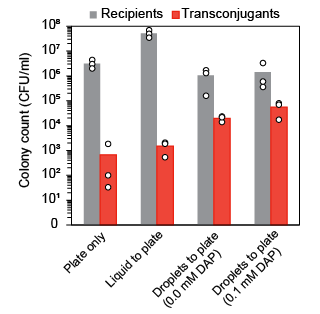

Figure S1. Recipient and transconjugant colony-forming units (CFU) used to calculate conjugation efficiency values shown in Figure 1. CFU were quantified by plating onto selective media for recipients and transconjugants, respectively. *S. venezuelae* recipients were selected by plating onto AS1 media supplemented with nalidixic acid to inhibit donor growth, while transconjugants were selected by the addition of apramycin to the media. Plates were incubated at 30 °C for 2-5 days and the resulting colonies were counted. Compared to cells incubated on plates, incubation in microdroplets produced between 10 and 100-fold increase in the absolute number of transconjugates.

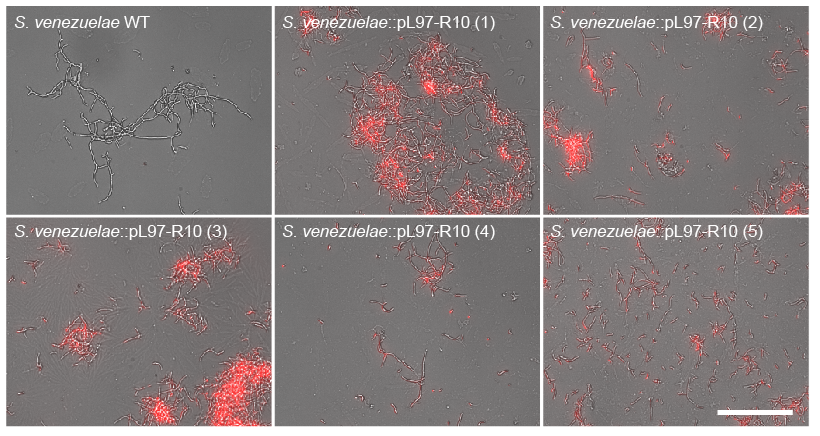

Figure S2. Fluorescence microscopy confirms mCherry expression in *S. venezuelae* transconjugants. Representative fluorescence microscopy image of wild-type *Streptomyces venezuelae* (WT) and five independently isolated transconjugant colonies. Following microdroplet-based conjugation transconjugant *S. venezuelae* colonies were imaged for mCherry fluorescence. Wild-type cells show no detectable red fluorescence, while transconjugants exhibit mCherry fluorescence, confirming successful plasmid transfer and expression. Scale bars, 100 µm.

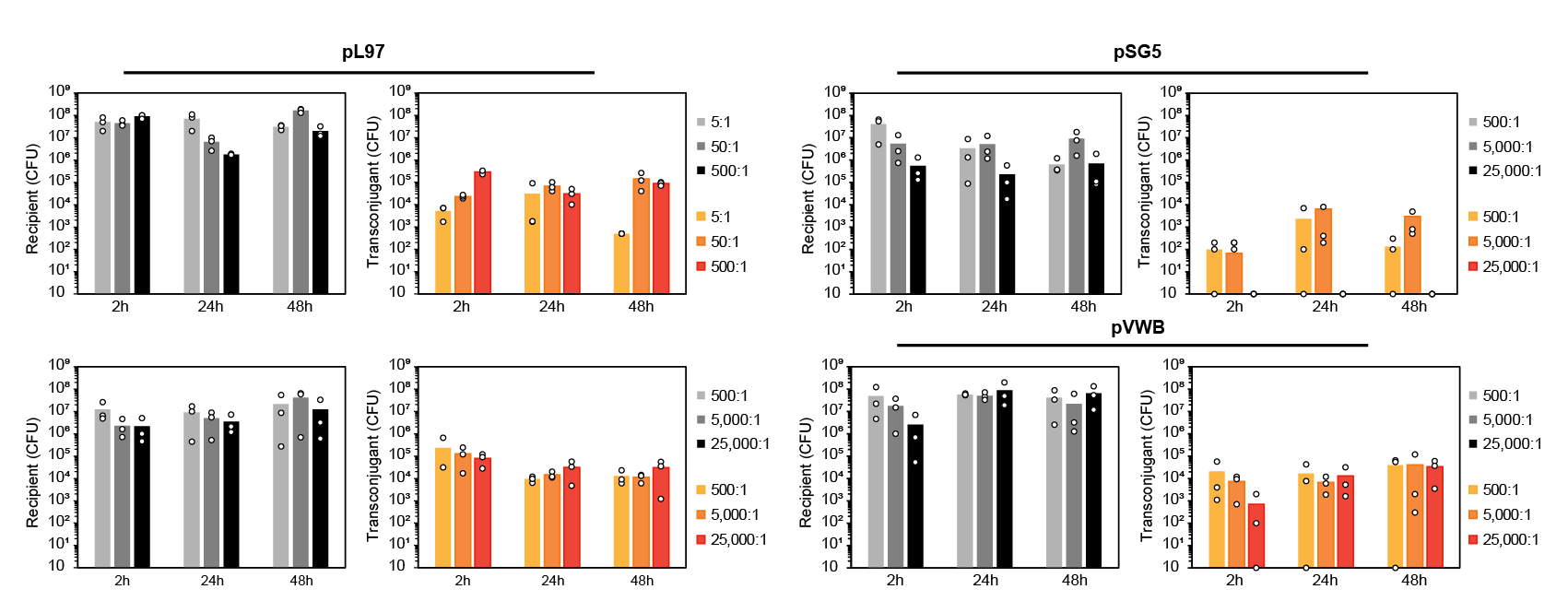
Figure S3. Recipient and transconjugant colony-forming units (CFU) used to calculate conjugation efficiency values shown in Figure 2. Recipient and transconjugant colony-forming units (CFU) were measured across varying donor to recipient ratios and incubation times in microdroplets. For pL97, donor-to-recipient ratios of 5:1, 50:1, 500:1, 5,000:1, and 25,000:1 were tested, while pSG5 and pVWB were only evaluated at ratios of 500:1, 5,000:1, and 25,000:1. Cells were incubated in microdroplets for 2 h, 24 h, or 48 h prior to plating. CFU were quantified by selective plating, with recipients selected on AS1 media supplemented with nalidixic acid and transconjugants selected with nalidixic acid and apramycin. Plates were incubated at 30 °C for 2–5 days before colony counting.

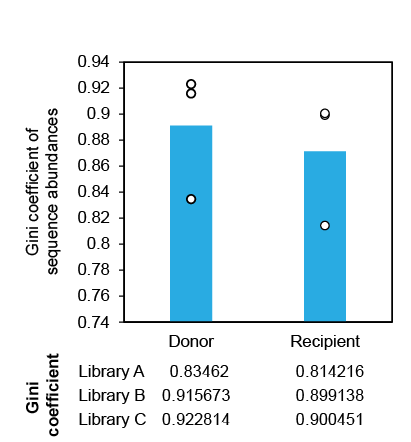

Figure S4. Gini coefficients of barcoded plasmid libraries. Comparison of barcode library uniformity before and after conjugation, quantified using the Gini coefficient of sequence abundances. Input libraries were sequenced from donor minipreps (Donor), and corresponding output libraries were recovered by PCR amplification from transconjugant *S. venezuelae* genome following microdroplet-based conjugation (Recipient). Each point represents an independently cloned library.

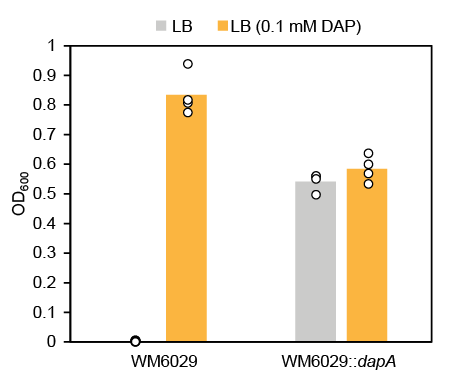

Figure S5. Evaluating *dapA* complementation on donor growth. *E. coli* WM6029 were transformed with a plasmid that either contained or lacked *dapA*. Following an overnight incubation at 37 °C, optical density at 600 nm (OD_600_) was measured. Cells that are not complemented with *dapA* are only able to grow in the presence of 0.1mM DAP, while in the absence of DAP, there was no difference in growth for the *dapA* complemented strain.

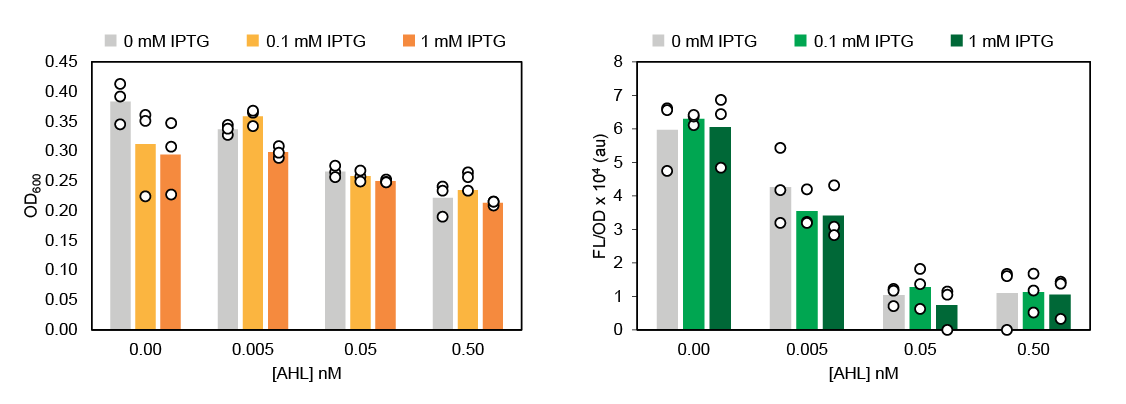
Figure S6. Suicide donor AHL/IPTG titration FL/OD raw values. To initially characterize AHL and IPTG induction for efficient donor removal via a holin and lysin mechanism, *E. coli* WM6026 was transformed with three plasmids: the two kill switch circuit plasmids and a conjugative plasmid that expresses dapA and sfGFP. Cells were grown in liquid culture overnight and subcultured into fresh media with varying concentrations of AHL (0 nM - 0.5 nM) and IPTG (0 mM - 1 mM). Cell density was measured after three hours of growth by optical density at 600 nm (OD600), and fluorescence characterization of the conjugative plasmid was performed (measured in units of fluorescence/optical density (FLOD).

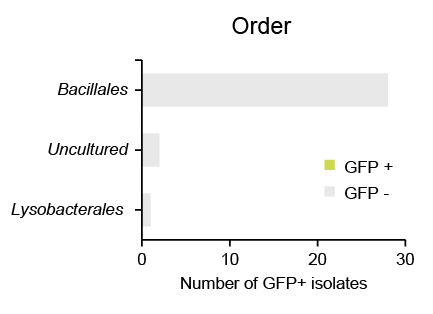

Figure S7. Colony PCR analysis of isolates from bulk community conjugation control. To assess plasmid transfer under conventional plate-based conjugation conditions and bulk liquid culture outgrowth, the soil microbial community was incubated with suicide donor cells on solid media without microdroplet encapsulation. After conjugation, cells were transferred to liquid media with AHL and IPTG to eliminate donor cells and allow transconjugant outgrowth. Following this growth period, cultures were serially diluted and plated, and 30 randomly selected isolates were screened by colony PCR for the presence of the GFP on the conjugative plasmid. No isolates tested positive for GFP, indicating a lack of detectable plasmid transfer. 16S sequencing of the isolates revealed that *Bacillales* were the single taxonomic order that was predominantly recovered, highlighting that bulk conjugation is both less efficient and less capable of preserving slower-growing or less competitive taxa.
